# Scouting ecological drivers of natural enemies in citrus orchards: implications for biological control in the Corsican agricultural landscape

**DOI:** 10.64898/2026.05.27.727191

**Authors:** Emma Carrié, Lucas Margris, Esther Carlut, Elise Frank, Elsa Canard, Manuel Plantegenest, Hervé Sanguin, Laurent Julhia, Virginie Ravigné, Samuel Soubeyrand

## Abstract

1. Effective pest control requires a better understanding of natural enemy ecology, particularly how their distribution responds to landscape structure and local management. Agricultural intensification has simplified landscapes, reducing biodiversity and constraining pest control. Landscape-scale surveys are therefore needed to identify strategies that support natural enemies in agricultural systems.
2. From 2021 to 2022, over three seasons, we surveyed 25 clementine orchards across a gradient of landscape complexity in Corsica. We modelled the tree-level occurrence of four natural enemy species in relation to ecological context across multiple spatial and temporal scales. Site-level evenness, reflecting differences in species’ occurrence frequencies, was also estimated to assess how natural enemy assemblages vary across local management and landscape gradients.
3. Climate and local management were the primary drivers of species occurrence, with additional contributions from landscape factors. Species displayed distinct responses, while model performance increased with the number of sampled trees, plateauing at around half of the sampling effort.
4. Site-level evenness was higher in organic orchards than in conventional orchards. In conventional orchards, it responded to landscape structure in a scale-dependent manner, increasing at the larger scale with the heterogeneity and spatial continuity of semi-natural habitats.
5. ***Synthesis and applications*.** Organic farming enhances natural enemy presence and diversity in Corsican citrus orchards, while the naturalness and complexity of surrounding landscape can mitigate species scarcity in conventional orchards. We recommend continued long-term monitoring of arthropod communities in these systems, prioritizing the number of sampling sites and years while reducing the number of trees sampled per site to optimize effort.

## 1. INTRODUCTION

In agricultural landscapes, arthropod communities provide key ecosystem services, including crop pollination and biological pest control, i.e. the suppression of pests by their natural enemies through antagonistic interactions like predation, parasitism or competition (McEvoy, 2018; Dainese et al., 2019). The increasing need to reduce pesticide use while maintaining crop yields makes reliance on biological pest control a promising strategy for sustainable agriculture. However, its effectiveness varies greatly over space and time. Understanding how environmental drivers and local management shape natural enemy populations is therefore essential to develop agroecological crop protection strategies and to anticipate pest outbreaks (Soubeyrand et al., 2024).

Conservation biological control promotes the survival and behavioral performance of natural enemies through targeted land-use management (Landis et al., 2020). Such efforts depend on the extent to which farm- and landscape-level characteristics influence populations of natural enemies. Landscape structure, often described as a gradient of landscape complexity, is a key determinant of biodiversity and biological pest control (Chaplin-Kramer et al., 2013; Haan et al., 2020). It comprises two interrelated components: composition, defined as the proportions of distinct land-use types (Lu et al., 2022; Plata et al., 2024), and configuration, referring to the size, shape, and spatial arrangement of land patches (Martin et al., 2019). The interactive effects of landscape composition and configuration vary according to the taxa involved, their functional traits, and the spatial and temporal scales at which they are examined (Martin et al., 2016; Martin et al., 2019). Nonetheless, empirical evidence generally supports the hypothesis that increasing landscape complexity, particularly through the presence of undisturbed habitats such as semi-natural vegetation, field margins, and hedgerows, enhances the diversity and abundance of natural enemies compared with structurally simple, intensively cultivated landscapes (Bianchi et al., 2006; Chaplin-Kramer et al., 2011; Lu et al., 2022). Multiple mechanisms are likely to operate simultaneously to underlie these patterns, including access to complementary food resources (e.g. alternative prey, pollen and nectar), overwintering sites, refuges from disturbance or intraguild predation, and increased habitat connectivity that facilitates movement and persistence (Landis et al., 2000; Langellotto and Denno, 2004). Most landscape-scale studies rely on structural connectivity metrics, based on the spatial attributes of land patches (e.g. Euclidean distances). While useful, these metrics do not account for species-specific responses. Functional connectivity, in contrast, integrates movement constraints imposed by habitat quality and barriers. Graph models can measure connectivity in relation to the spatial configuration of the patches, the dispersal capacities and ecological requirements of species (Rayfield et al., 2011; Tarabon et al., 2021).

Local farming practices can interact with these landscape mechanisms. Organic farming (e.g. avoiding synthetic pesticides, herbicides, mineral fertilisers, natural enemies input) generally enhances arthropod abundance and diversity within fields (Bengtson et al., 2005; Winqvist et al., 2011; Martin et al., 2016). Its effectiveness for pest control depends on the surrounding landscape structure and remains debated (Birkhofer et al., 2016; Muneret et al., 2018; Laffon et al., 2024). Most studies addressing how landscape structure and local management affect pests and natural enemies concern annual crops, with comparatively few examples in perennial systems (Etienne et al., 2024; Laffon et al., 2024; Plata et al., 2024; Romero et al., 2025). The permanence, multi-strata structure and hedgerows typical of orchards can support stable natural-enemy communities (Simon et al., 2010, Demestihas et al., 2017, Herz et al., 2019), whereas intensive pesticide use may disrupt them. Overall, predicting the ecological contexts that maximize pest control remains challenging (Alexandridis et al., 2021; Bonato et al., 2023). This highlights the need for a better understanding of the local ecological signals shaping natural enemy assemblages within particular agroecosystems.

Here, we first investigated the relative importance of ecological and local management factors on natural enemy occurrence within a landscape monitoring network across citrus orchards in Corsica (France), a major economic and cultural cropping system. In citrus orchards, natural enemy communities include both native and regularly introduced species used for scale-insect control, forming a largely understudied system. We focused on four key natural enemies: the green lacewing, *Chrysoperla carnea*, and the three Coccinellidae, *Harmonia axyridis*, *Exochomus quadripustulatus*, and *Cryptolaemus montrouzieri*, which are important for the biological control of hemipteran pests (Bibi et al., 2026). These species were selected as potential predators of psyllid vectors of Huanglongbing (Vandenberg et al., 1987; Ruíz-Rivero et al., 2021). We applied a model-fitting and -selection approach to examine species-specific responses to local- and landscape-scale factors. This approach was also leveraged to quantify the information content in size-increasing sampling designs about the link between natural enemy occurrence and factors, in the aim of allocating future monitoring efforts in a rational way. We also assessed natural enemy evenness at the orchard level and examined its relationship with local management and landscape structure. We hypothesized that organic farming would enhance natural enemy occurrence and assemblage diversity, thereby potentially reducing pest pressure, especially in simple, intensively cultivated landscapes in comparison with more complex landscapes that include semi-natural habitats.

## 2. MATERIAL AND METHODS

### 2.1 Study system and sampling design

To document the influence of the agricultural system (organic *vs* conventional) and the landscape structure on the presence of natural enemies in citrus orchards, we sampled 25 clementine (*Citrus clementina*) orchards in eastern Corsica (Fig. 1), the main citrus producing region of the island. Orchards were selected along a gradient of semi-natural habitat proportion within their surrounding landscape (radius 500 m). The survey encompassed both juvenile (< 5 years old) and mature (> 15 years old) orchards, reflecting the marked age heterogeneity of citrus orchards in this region. Orchards were chosen so as to examine an effect of local management, across conventional (n = 12) and organic (n = 13) production systems. A fine characterization of practices was not available, although management intensity in Corsican citrus orchards is relatively low compared with other crops (Agreste, 2021). Average orchard area was 5.0 ± 2.5 ha for organic and 5.0 ± 4.7 ha for conventional orchards (mean ± SD).

**FIGURE 1.**
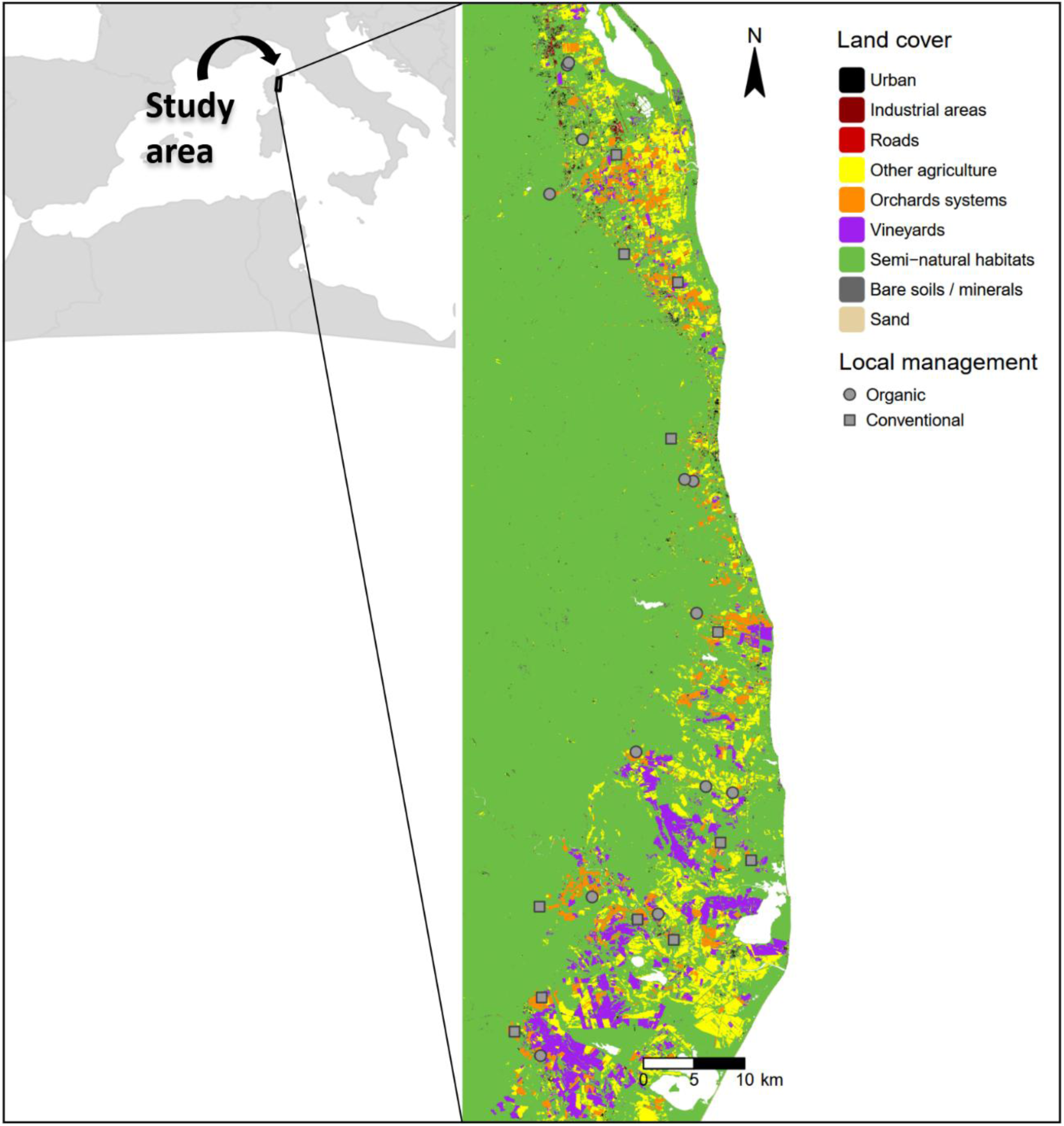
Map showing the locations of orchards selected for this study in eastern Corsica, with land cover and local management types indicated.

### 2.2 Arthropod Sampling

To assess natural enemy occurrence and its spatial and temporal variation, 40 trees were sampled in each orchard over two consecutive years (2021 and 2022) and three seasons: April/May, July/August, and September/November. We used a standardized back-and-forth sampling transect across the plot (Appendix S1, Fig. S1). The transect comprised 10 evenly-spaced sampling stops, and at each stop arthropods were surveyed on two adjacent trees on two adjacent rows, resulting in a total of 40 sampled trees per orchard. Target arthropod species were sampled by beating branches over a white beating sheet. Each branch was struck three times, and all dislodged individuals were collected and subsequently identified in the laboratory. A circular hoop (1 m diameter) was also randomly positioned in front of each tree to delineate a standardized sampling area, within which flushes and flushes infested by mealybugs (*Planococcus citri*) and leafminers (*Phyllocnistis citrella*) were counted. Flush infestation data were used to quantify tree-level pest pressure, although related analyses were beyond the scope of the present study and are therefore provided in Appendix S2.

For the subsequent analyses, we focused on four predators: (i) the green lacewing (*C. carnea*), a generalist predator and native biological control agent in orchards, relatively tolerant to insecticides (Porcel et al., 2013; Paredes et al., 2024); (ii) the pine ladybird (*E. quadripustulatus*), a generalist predator sometimes released in Corsican citrus orchards (Holecová et al., 2018; Di Sora et al., 2024); (iii) the mealybug destroyer (*C. montrouzieri*), a ladybird native to Australia and one of the most widely released agents for mealybug control in citrus worldwide, including Corsica (Ferreira et al., 2021; Moretti et al., 2025); (iv) the harlequin ladybird (*H. axyridis*), an invasive Asian ladybird and generalist predator capable of intraguild predation on other ladybirds, with high adaptability to diverse habitats including urban areas (Vandereycken et al., 2012; Honek et al., 2021; Collop et al., 2024).

### 2.3 Climate and soil data

Climate was characterized using 19 bioclimatic variables (O’Donnell and Ignizio, 2016) derived from the SAFRAN model (Météo France), which interpolates daily observations from over 1000 meteorological stations at 8-km resolution. SAFRAN climatic data were downloaded via the SICLIMA platform developed by AgroClim-INRAE. Bioclimatic indices for 2021 and 2022 were computed separately for each year, based on monthly minima, maxima, and mean air temperature (2-m above ground) or precipitation, to capture spatial and interannual climatic variability. In addition, seasonal weather variables for spring (March–May), summer (June–August), and autumn (September–November) were obtained by averaging daily values of temperature (mean, minimum and maximum), precipitation, runoff, relative and specific humidity, actual and potential evapotranspiration, visible radiation and wind speed. Site-level soil chemical elements, topography and land orientation were retrieved from Portes et al. (2024), with all multisource data standardized to a 500 m × 500 m spatial resolution grid. Detailed descriptions of the climatic, soil and topographic variables are provided in Appendix S3, Table S2.

### 2.4 Landscape characterisation

To explore the effects of landscape structure on the focal natural enemy species, we extracted land cover information surrounding each orchard at multiple spatial scales. “Scale” is defined in this study as the buffer radius around the survey area within which landscape variables were calculated. Concentric buffers of 500, 1500, and 3000 m were created from the outer edge of the survey area. These radii align with the typical dispersal distances of the focal species (Table S3 in Appendix S4) and on previously reported landscape effects on arthropod diversity (Paredes et al., 2024; Plata et al., 2024; Romero et al., 2025). Maps were created by combining: (i) the OSO Land Cover 2022 product developed by the Land Cover Scientific Expertise Centre of the French Theia Land Data and Services Centre, which provides a 10-meter resolution raster land-cover classification (24 classes) derived from Sentinel-2 time series; (ii) the regional growers’ organisation (AOP Fruit de Corse) citrus parcel dataset; and (iii) the full road network from OpenStreetMap. First, citrus polygons were rasterized and merged into the original OSO land cover map, creating a new subclass of the ‘orchard’ class (class 14) for citrus. Secondly, the road network was extracted from OpenStreetMap using the R packages ‘osmdata’ (Padgham et al., 2017) and ‘sf’ (Pebesma, 2018) to complement the ‘road’ class of the OSO raster. Three original OSO categories were then grouped into the new class for semi-natural habitats (SNH), which included forests (broadleaf and coniferous) and shrublands. Several compositional and configurational variables were calculated for the buffer zone using the R package ‘landscapemetrics’ (Hesselbarth et al., 2019), at the landscape and class level. For the class-level variables, we focused on functionally-important land covers in the study area: citrus orchards, SNH, vineyards, urban areas and roads. Definitions for all landscape metrics are provided in the package documentation (Hesselbarth et al., 2019), while Table 1 lists the variables reported in the present study.

**TABLE 1.**
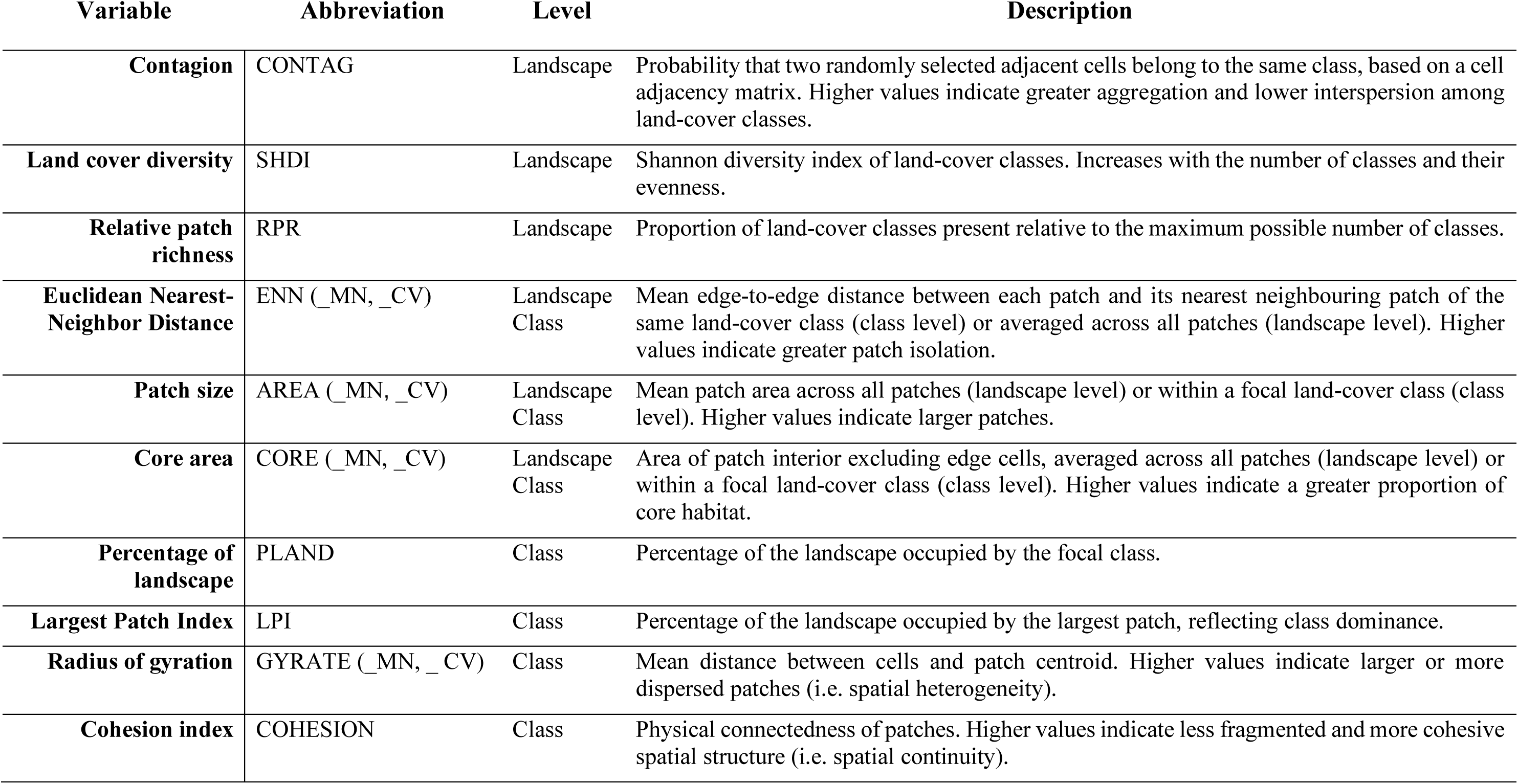
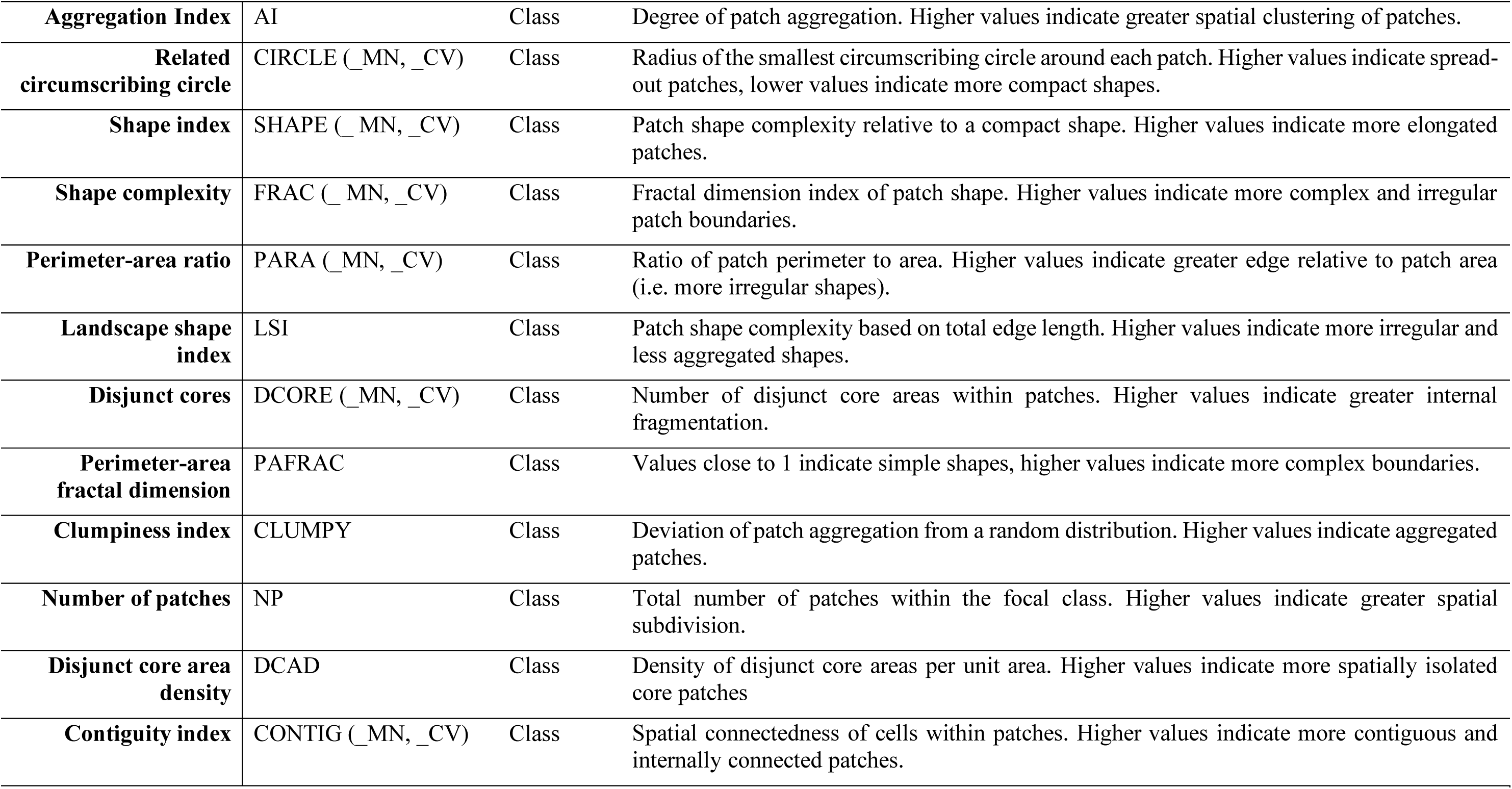
List and description of the landscape metrics reported in this study. Metrics were calculated from a 10-m resolution land cover raster using the R package ‘landscapemetrics’ (v2.2.1), at the class or landscape level. Suffixes in variables abbreviations: _MN denotes mean values, _CV denotes coefficient of variation, where applicable.

### 2.5 Functional connectivity

The R package ‘graph4lg’ (Savary et al., 2020), which integrates Graphab functionalities (Foltête et al., 2021), was used to model habitat networks at the 3000 m scale. Citrus orchards were defined as habitat patches (nodes) and linked through least-cost paths derived from a resistance surface. To build the resistance surface, land use categories were assigned to one of five levels of resistance (Morin et al., 2024), according to species’ ecology (Appendix S4, Table S3). We calculated a global metric, namely the Probability of Connectivity index (PC; Saura and Pascual-Hortal, 2007), defined as ‘the probability that two propagules randomly placed within the landscape fall into habitat areas that are reachable from each other’:

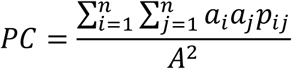

where *a_i_* and *a_j_* are the area of patches *i* and *j*, *A* is the total area of the study zone and *p_ij_* is the probability of dispersal between patches *i* and *j*. Dispersal probability was modeled as an exponential decay function of least cost distance: *p_ij_* = exp(-α*d_ij_*), where *d_ij_* is the least-cost distance between patches and α controls the decay rate. Following Saura and Pascual-Hortal (2007), the decay parameter α was calibrated so that dispersal probability equals 0.05 at the maximum dispersal distance defined from the literature (Appendix S4, Table S3).

We also computed a local metric for each orchard: a centrality metric based on circuit theory (Current Flow CF, Girardet et al., 2015). Links (between habitat patches in the network) were treated as resistors and habitat patches as current sources. The current flowing through the network represents potential dispersal flux. Patch capacity was assumed proportional to patch area (β = 1).

### 2.6 Modelling

We implemented a multi-step framework to identify, quantify, and validate the key drivers of natural enemy temporal and spatial distributions and assemblages (Fig. 2). Data curation and statistical analyses were conducted using R (R Core Team, 2022). Observed abundances were generally low and most sites contained 0 or 1 individual per species at the tree scale. Therefore, all subsequent analyses were performed using occurrence data.

**FIGURE 2.**
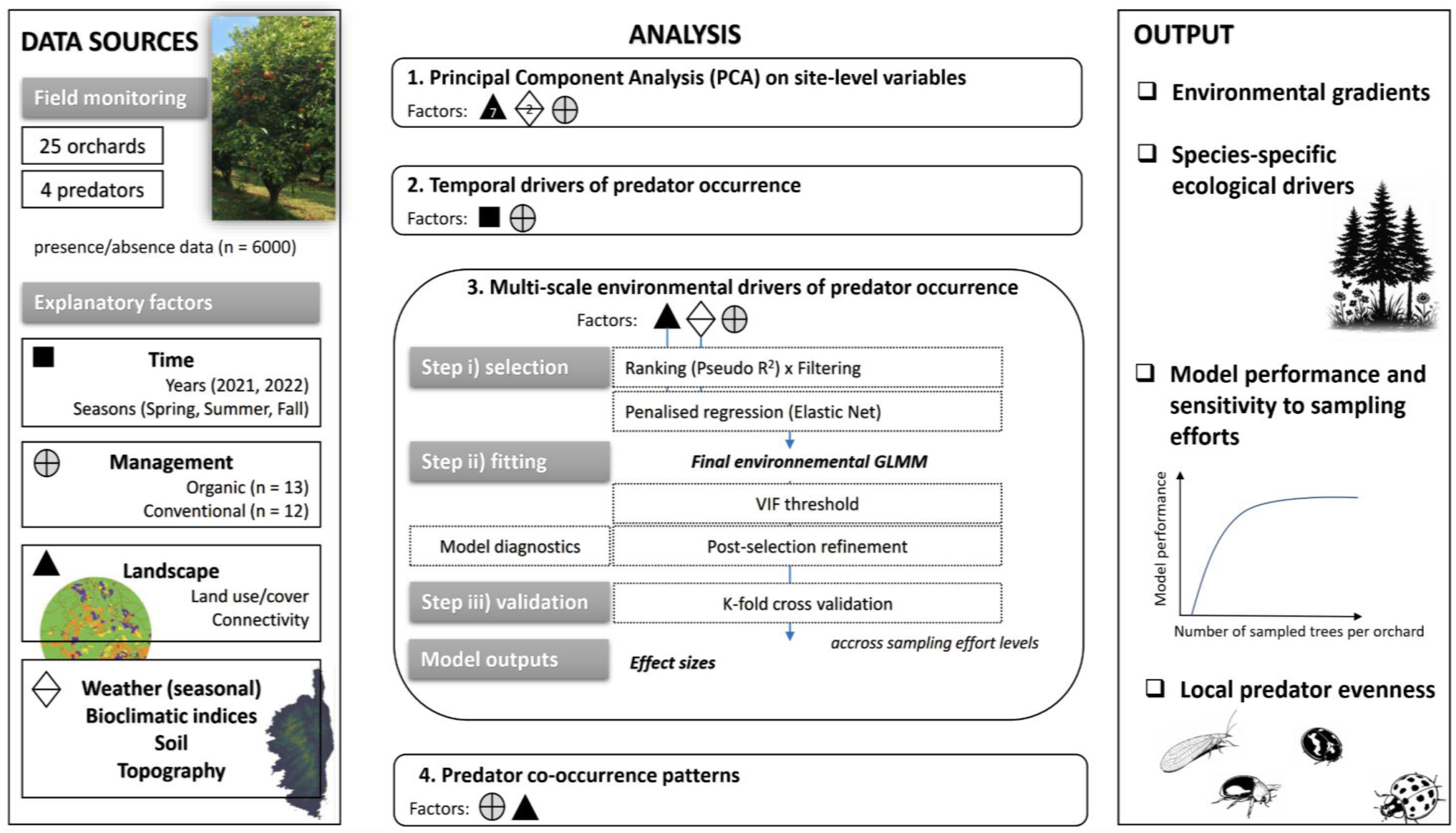
Workflow of field data collection and analyses (Principal Component Analysis and binomial Generalized Linear Mixed Models) used to identify ecological drivers of natural enemy occurrence and evenness across local management types in the Corsican agricultural landscape. Symbols indicate groups of explanatory variables used in the analyses: time (black square), local management (cross in a circle), landscape (black triangle), and other environmental conditions (white diamond).

#### 2.6.1 Landscape and environmental variation across plots and scales

First, to identify major regional landscape gradients and the distribution of organic and conventional orchards along them, we performed a principal component analysis (PCA) including a subset of seven variables capturing key aspects of landscape complexity within 1500 m buffers. Landscape aggregation (CONTAGION) and land cover diversity (SHDI) were included to represent configurational and compositional heterogeneity (based on Galpern et al., 2021). Class-level dominance variables (LPI) for citrus, SNH, and urban land covers, as well as site elevation and annual mean temperature (BIO1, for 2021) were included as descriptors of the ecological context. The PCA was performed with the PCA() function in the R package ‘FactoMineR’ (Lê et al., 2008).

Second, we focused on semi-natural habitat (SNH) given their strong hypothesized functional role for natural enemies and the significance of some configurational metrics in our models. Pairwise Pearson correlations were assessed for composition (PLAND) and configuration metrics (COHESION and GYRATE_CV, Table 1) describing SNH at two contrasted spatial scales (500 and 3000 m). This approach allowed us to assess how the relationships between composition and configuration, and among configuration metrics themselves, vary across spatial scales. These analyses are important for interpretation because, while SNH is generally expected to support natural enemy populations, the effects of configurational features are often ambiguous and highly scale-dependent (Zhang et al., 2020). Significance of correlations was tested using the cor.mtest() function and visualized with the ‘corrplot’ R package (Wei and Simko, 2021).

#### 2.6.2 Temporal variation in natural enemy occurrence across management types

We examined temporal variation in natural enemy occurrence across local management types. The analysis was carried out at the tree scale, where the response variable was binary, i.e. corresponding to whether or not the natural enemy was observed. Binomial Generalized Linear Mixed Models (GLMMs) followed by a type-III test of deviance (or type-II when no interaction was included), were used to assess the effects of the year, season and local management on the probability of occurrence. Orchard was included as a random effect to account for the grouping structure of the data. A first analysis was performed with year as a fixed effect factor. Subsequently, for each year separately, we performed a second analysis with season, local management and their interaction as fixed factors.

#### 2.6.3 Environmental drivers of natural enemy occurrence

We assessed the relative importance of climatic, soil, topographic and landscape characteristics on natural enemy occurrence. Factors were initially ranked by their univariate association (marginal pseudo-R²) with the binary response variable. For each pair of factors with Pearson correlation above 0.75, the lower-ranked factor was removed, retaining the one most strongly associated with the response, in order to minimize redundancy (Chauvin et al., 2025). Relevant factors were then selected using a shrinkage approach (Elastic Net; Zou and Hastie 2005). The method combines Lasso and Ridge penalties, shrinking coefficients of less important factors toward zero. Optimal values of the tuning parameters (α, λ) were selected using cross-validation with the ensr() function (DeWitt & Bennett, 2019). Factors with non-zero coefficients were retained and added to a naive model, which included orchard and season nested within year as random effects, and local management as a fixed effect. Collinearity was assessed using the variance inflation factor (VIF). Factors with VIF>3 were removed iteratively until all remaining factors had VIF<3. All factors were scaled, allowing comparison of effect sizes in the model. A post-selection refinement step evaluated the structural importance of the least influential predictors based on absolute β coefficients. Individual and combined removal had a negligible effect on model AUC (<1%), justifying their exclusion to simplify the model. Subsequently, two-way interactions between local management and landscape factors were tested. Spatial autocorrelation was tested with Moran’s I (none detected), and residuals were inspected with the R package ‘DHARMa’ (Hartig, 2022) to check for overdispersing.

Model cross-validation was performed using a 10-fold grouped design, with folds defined by unique combinations of site, year, and season. Entire groups were held out from model fitting to prevent data leakage between training and testing sets. Within each training fold, trees were randomly subsampled within groups to assess the effect of sampling effort on model performance. This procedure was repeated in 600 cross-validation runs for each sampling scenario. For each run, binomial GLMM performance was evaluated using AUC and balanced accuracy. Classification thresholds were optimized by maximizing the mean of sensitivity and specificity. Additional validation analyses assessing model performance under site level cross-validation are provided in Appendix S5.

#### 2.6.4 Site-level evenness across management types and landscape structure

The joint occurrence of the natural enemy species was approached by estimating their diversity using the R package ‘iNEXT’ (Hsieh et al., 2016), on incidence data (i.e. presence/absence of species) from all collection events at each site, defined as individual trees surveyed during each season and year. Hill numbers of order q = 1 (the exponential of Shannon entropy, Exp(*H′*)) were calculated. Accumulation curves were generated, and site-level Hill numbers were extracted. These estimates were subsequently used to explore correlations with the landscape factors. Sample completeness was assessed via coverage-based rarefaction and extrapolation to evaluate whether the sampling effort was sufficient. Confidence intervals (95%) were computed from 1000 bootstrap replicates.

GLMMs were fitted using the ‘lme4’ package (Bates et al., 2015). A random effect for the tree itself was always tested but removed, as it did not significantly improve model fit. Step-by-step top-down strategy was used to test the two-ways interaction terms, and non-significant interaction terms were removed. Multiple post-hoc comparisons of estimated marginal means were conducted with false discovery rate (‘fdr’) P-value adjustment method, on significant interaction effects or significant main effect(s) when no interaction term was significant.

## 3. RESULTS

The landscape composition within a 500 m radius of the monitored orchards varied considerably: semi-natural habitats, “SNH” (26.8–91.2%); citrus orchards (2.2–27.2%) and vineyards (0–37.8%).

### 3.1 Structural gradient in the Corsican agricultural landscape

The first two dimensions of the PCA explained 59.3% and 19.2%, respectively, of the total variability among orchards with respect to environmental and landscape input factors (Fig. 3a). The first dimension expressed mostly the variability in landscape aggregation (CONTAGION), land cover diversity (SHDI), SNH dominance (LPI), annual temperature (BIO1), and elevation. The second dimension expressed mostly the variability in citrus and urban dominance, as well as elevation. Overall, it revealed a gradient ranging from higher-elevation sites embedded within large SNH to coastal zones dominated by mixed farming interspersed with urban areas. Hotelling’s test indicated no significant difference in orchard positions between organic and conventional systems, consistent with the overlap of their 75% confidence ellipses (Fig. 3a).

**FIGURE 3.**
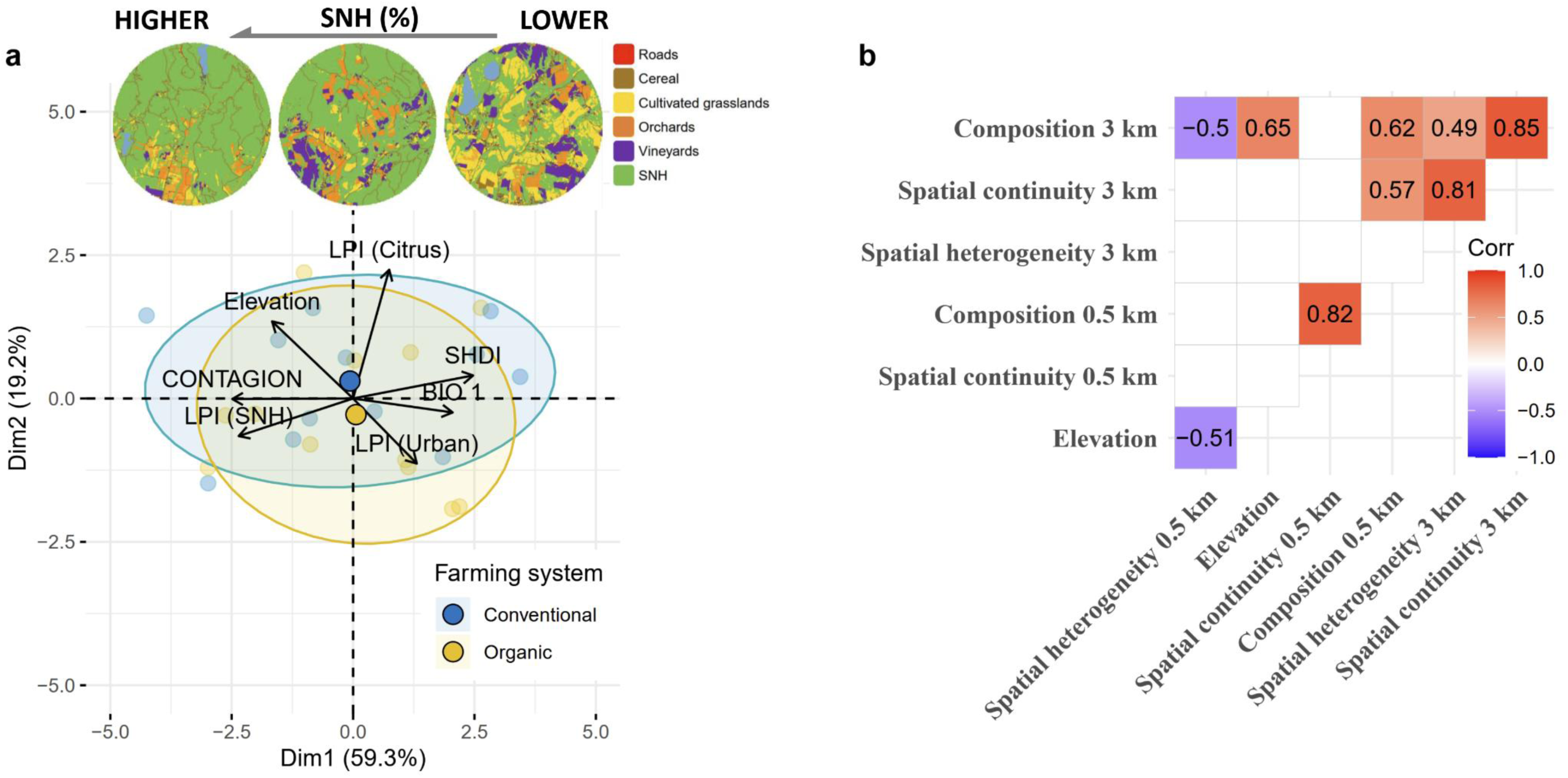
PCA biplot showing the relationship between landscape (buffer of 1500 m radius) and environmental variables, as well as the distribution of organic and conventional orchards along the first two PCA dimensions, including barycenters and 75%-confidence ellipses for local management types (a). Metric abbreviations are as follows: CONTAGION (landscape aggregation), SHDI (land cover diversity), LPI (largest patch index for specific land-cover classes, including citrus orchards, urban areas, and semi-natural habitats, ‘SNH’), and BIO1 (mean annual temperature). Pearson correlation coefficients (only significant values shown) among some SNH metrics: spatial heterogeneity (GYRATE_CV), spatial continuity (COHESION), and composition (PLAND) at 500 and 3000 m scales (b). Buffer radii are expressed in km in panel (b) for readability.

Pairwise Pearson correlations revealed relationships among compositional (PLAND) and configurational components of SNH, including spatial heterogeneity (GYRATE_CV) and continuity (COHESION) (Fig. 3b). For each scale (at 500 and 3000 m), SNH composition and spatial continuity were strongly correlated. In contrast, spatial heterogeneity was only moderately correlated with composition, and only at the larger scale (r = 0.49). Similarly, configurational heterogeneity was highly correlated with continuity, but only at the largest scale (r = 0.81). Cross-scale correlations showed that SNH composition at local and large scales was moderately related (r = 0.62), whereas configurational metrics exhibited no significant cross-scale correlations, supporting the relevance of our multi-scale analysis.

### 3.2 Temporal patterns of natural enemy occurrence across management types

The occurrence probability of *C. carnea* (Fig. 4a) varied significantly between years (*P* < 0.001), with mean values higher in 2022 than 2021. In 2021, occurrence was affected by local management (*P* < 0.05) not by season, with higher mean values in conventional than organic orchards. In 2022, occurrence was affected by season (*P* < 0.001) and local management (*P* < .05). It was higher in spring compared with summer or autumn, and remained higher in conventional than in organic orchards.

**FIGURE 4.**
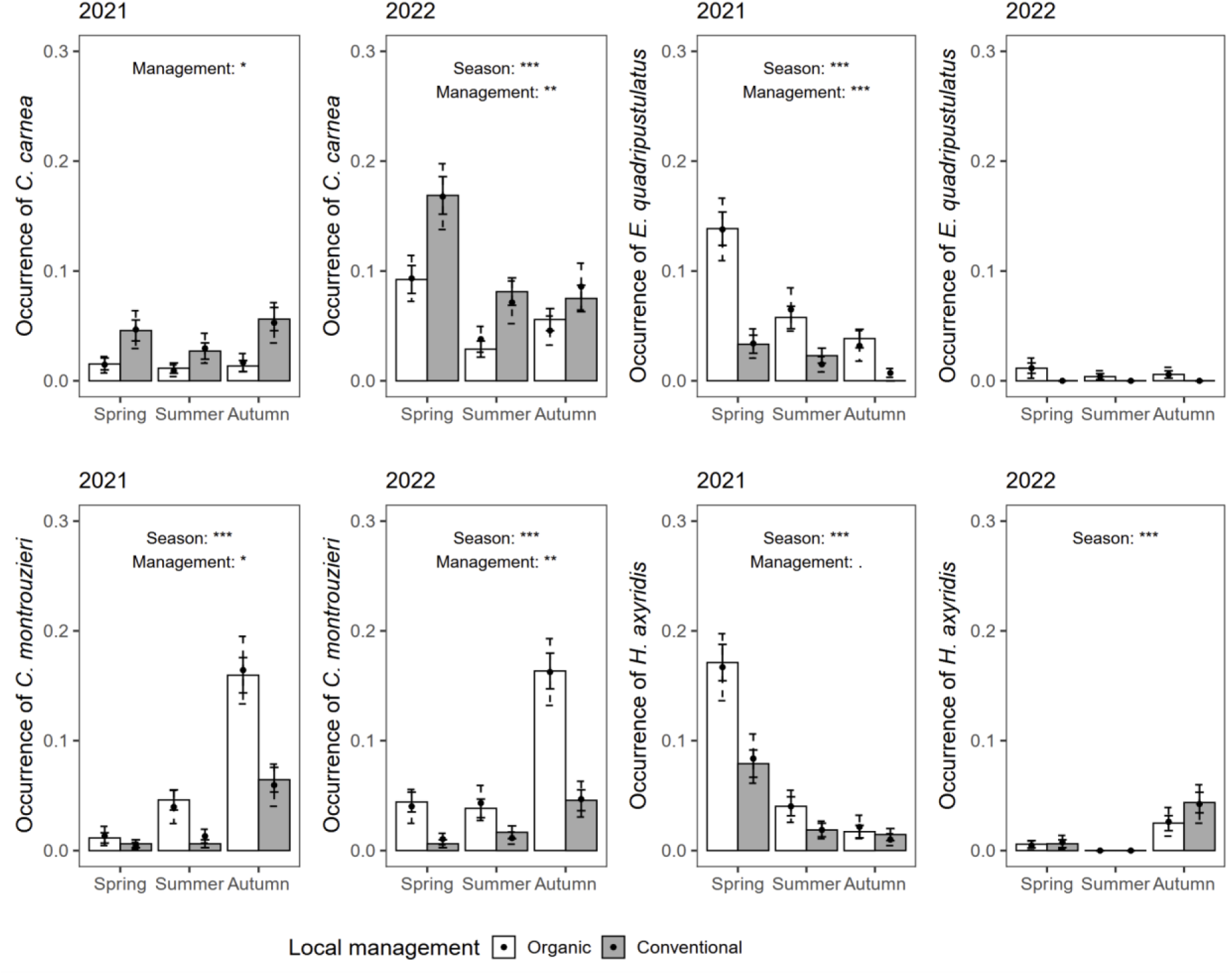
Probability of occurrence of *C. carnea* (a), *E. quadripustulatus* (b), *C. montrouzieri* (c) and *H. axyridis* (d), according to the year, season and local management type. Color bars and solid error bars indicate observed probabilities and 95% confidence intervals, whereas black points and dashed error bars indicate model estimates and their corresponding confidence intervals. For each year and species, significance of main effects (season, management) is indicated within panels. No significant interaction between season and local management was detected. Significance levels: *P* < 0.05 (*), *P* < 0.01 (**), *P* < 0.001 (***).

Occurrence of *E. quadripustulatus* (Fig. 4b) differed significantly between years (*P* < 0.001), with mean values higher in 2021 than 2022. In 2021, occurrence was affected by both season and local management (*P* < 0.001). It decreased from spring to summer and autumn, and was consistently higher in organic than in conventional orchards. In 2022, occurrence was not affected by either the season or local management.

Occurrence of *C. montrouzieri* (Fig. 4c) was not affected by the year. In both years, occurrence was affected by the season (*P* < 0.001) and local management (*P* < 0.05; *P* < 0.001). It was on average higher in autumn compared to summer and spring, and consistently higher in organic than in conventional orchards.

Occurrence of *H. axyridis* (Fig. 4d) varied significantly between years (*P* < 0.001), with mean values higher in 2021 than 2022. In 2021, occurrence was affected by season (*P* < 0.001) but not by local management (*P* = 0.07). It decreased from spring, to summer and autumn. In 2022, occurrence was only affected by seasons (*P* < 0.001), with higher mean values in autumn than summer or spring. The interaction terms were not significant.

### 3.3 Species-specific ecological drivers

Model performance increased with sampling effort (Left panels, Fig. 5) and stabilized after sampling 14–20 trees (mean ± SD: 18 ± 3.5) for *C. carnea*, *E. quadripustulatus* and *C. montrouzieri*, exceeding that of the naive model after sampling a small number of trees (<5). The model for *H. axyridis* only consistently outperformed the naive model when all climatic variables were excluded and more than about 15 trees were sampled (Fig. 5d), although predictive performance remained weak for this species (<0.7).

**FIGURE 5.**
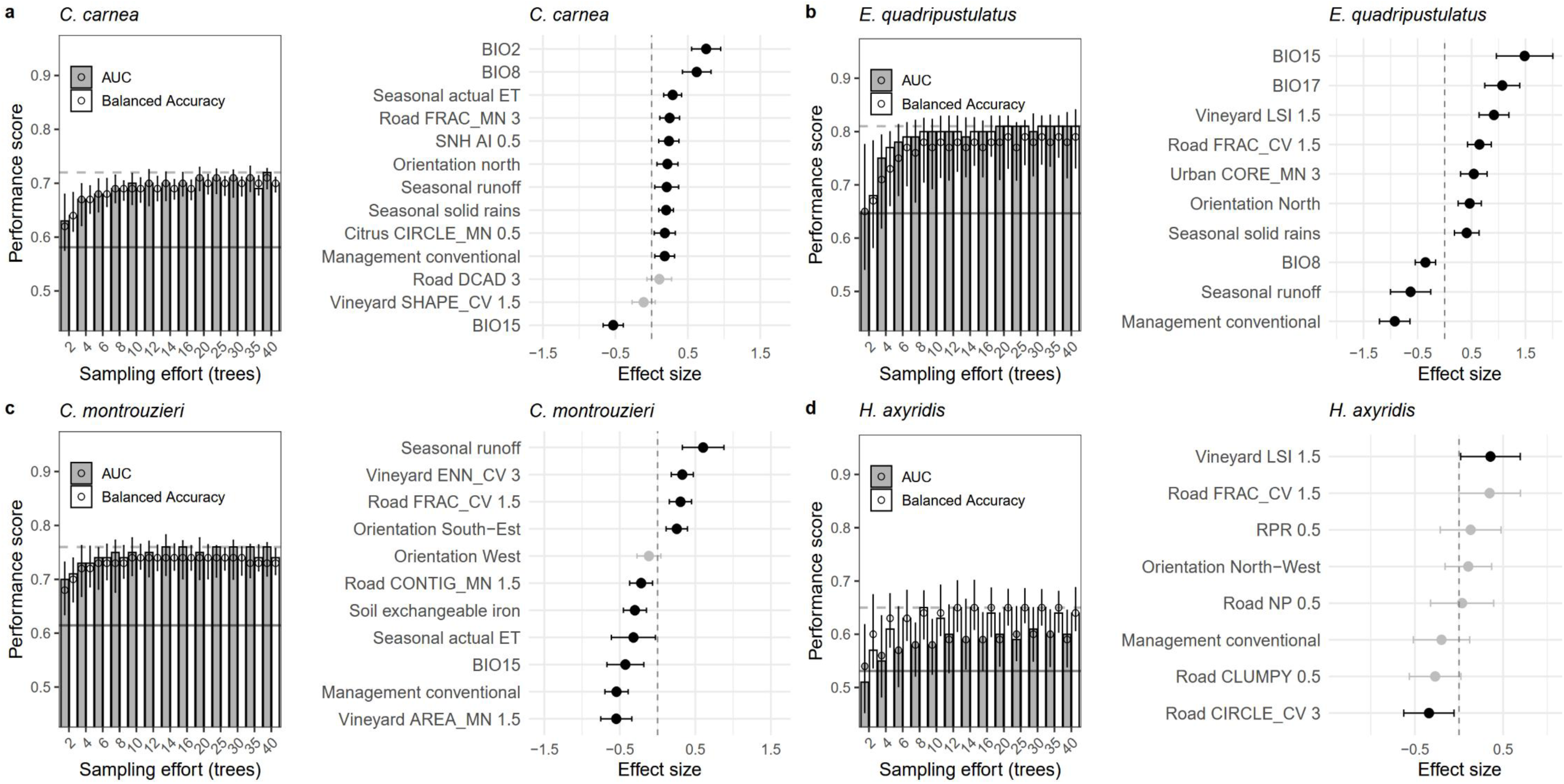
Left panels. Predictive performance of the binomial generalized linear mixed-effects model (GLMM) evaluated by cross-validation as a function of the sampling effort, i.e. the number of trees in the calibration dataset. Bars are median values across 600 repetitions, while points indicate mean estimates with their 95% confidence intervals. The plain horizontal line indicates the performance of the naïve model including only random effects and local management. The dotted horizontal line indicates the maximum performance across sampling scenarios. Right panels. Standardized effect sizes of the selected explanatory factors on natural enemy occurrence at the tree scale, derived from the GLMM. All continuous variables were scaled prior to analysis. For clarity, landscape variables include the spatial scale of calculation (buffer radii), expressed in km (0.5, 1.5 and 3 km), and are named as [land-cover class] [metric] [spatial scale]. SNH: semi-natural habitats. BIO variables correspond to bioclimatic variables (see Appendix S3 for detailed definition). Seasonal actual ET refers to actual evapotranspiration (mm). The coefficient associated with local management is not directly comparable to those of continuous predictors, as it represents a categorical factor. Black filled circles denote statistically significant effects (*P* < 0.05), whereas grey filled circles indicate non-significant effects.

Climatic variables showed the strongest associations with natural enemy occurrence, although responses were species-specific (Right panels, Fig. 5). Occurrence of *C. carnea* increased with diurnal temperature range (BIO2) or temperature of wettest quarter (BIO8) and decreased with precipitation seasonality (BIO15). Occurrence of *E. quadripustulatus* was positively associated with BIO15 and precipitation of the driest quarter (BIO17), while seasonal runoff (mm) had a negative effect. For *C. montrouzieri*, seasonal runoff had the strongest positive effect on the predator occurrence.

Landscape structure further shaped natural enemy presence, with no interaction with local management. Depending on the species, positive effects were associated with class-level configuration metrics, including SNH aggregation (AI, 500 m), road complexity (FRAC_MN, 3000 m; FRAC_CV, 1500 m), citrus patch compactness (CIRCLE_MN, 500 m), urban core area (CORE_MN, 3000 m), vineyard shape irregularity (LSI, 1500 m) and variation in inter-patch distance (ENN_CV, 3000 m). Negative effects were observed for vineyard patch size (AREA_MN, 1500 m), road contiguity (CONTIG_MN, 1500 m) and variation in patch compactness (CIRCLE_CV, 3000 m).

### 3.4 Site-level evenness across local management types and landscape structure

Rarefaction and extrapolation curves for site-level Hill diversity (q = 1) are shown in Fig. 6a. Because natural enemy assemblages comprised only four species, variation in diversity (q = 1) largely reflects differences in species’ occurrence frequencies rather than variation in richness, and can therefore be interpreted as evenness (Appendix S6). Sample completeness ranged from 0.85 to 1.00, indicating that the sampling effort was sufficient to reliably estimate diversity. Evenness differed significantly between local management types, being lower in conventional than in organic orchards (Wilcoxon test, *P* < 0.05; Fig. 6b), corresponding on average to assemblages of 2.7 versus 3.3 equally frequent species, respectively. Evenness was unrelated to landscape variables in organic orchards but significantly associated with 18 variables in conventional orchards (Fig. 6c, and illustrated in Appendix S6). These landscape features influencing site-level evenness shifted with spatial scale. At the largest scale (3000 m), evenness was positively correlated with SNH spatial continuity or heterogeneity (COHESION, GYRATE_CV), and vineyard complexity (PAFRAC). Conversely, citrus orchards spatial continuity, vineyard core area (CORE_MN) and the mean fractal dimension of SNH patches (FRAC_MN), were negatively correlated with evenness. At intermediate and fine scales, citrus variables at 1500 m were overall negatively correlated with diversity (except patch contiguity), just as road variables at 500 m (except patch contiguity).

**FIGURE 6.**
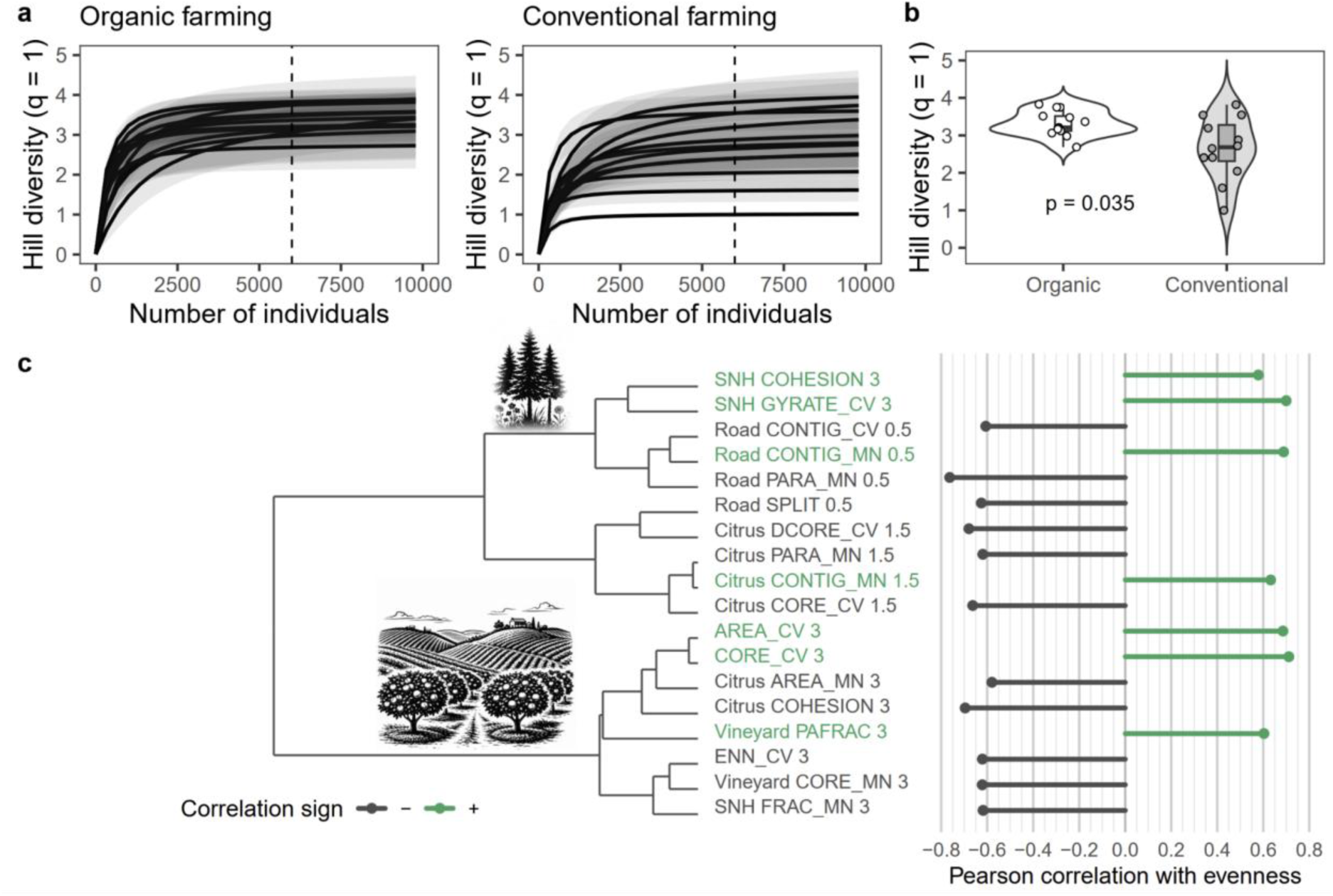
Rarefaction and extrapolation curves of natural enemy assemblages as a function of sampling units, based on Hill numbers with q = 1, shown separately for organic and conventional orchards (a). Vertical dashed lines indicate the cutoff between observed (rarefaction) and extrapolated portions of the curves. Each orchard is represented by a separate curve; shaded areas indicate 95% confidence intervals obtained by bootstrap resampling (1000 iterations). Comparison of site-level evenness between organic and conventional orchards using a Wilcoxon rank-sum test (b). Relationships between landscape variables and evenness in conventional orchards (c). Landscape variables are clustered based on their similarity (dendrogram, left), while horizontal bars show the direction and strength of Pearson correlations with evenness (right). Landscape variables include the spatial scale, expressed in km (0.5, 1.5 and 3 km), and are named as [land-cover class] [metric] [spatial scale].

## 3. DISCUSSION

The present landscape-scale experimental study in eastern Corsica highlighted the pattern of occurrences and its variations over time and space of four natural enemies and showed that their responses were jointly shaped by local management and landscape structure. We found partial support for our initial hypothesis: organic management increased site-level evenness, whereas its effects on natural enemy occurrence varied among species and, to a lesser extent, across seasons. This context dependency highlights that the benefits of organic practices do not translate uniformly into effective pest suppression. By integrating heterogeneous ecological data across scales, we identified the main drivers of natural enemy distribution and defined the sampling effort requirements for future monitoring strategies.

The contrasted patterns of occurrences observed among species reflect specific responses to climatic conditions, particularly temperature and humidity, which are key drivers of arthropod phenology and activity (Buckley et al., 2017; Grecchi et al., 2025). Notably, the autumn peak observed for *C. montrouzieri* is consistent with the seasonal increase in mealybug abundance in Mediterranean citrus orchards (Plata et al., 2024), reflecting the predator–prey population dynamics of this species (Pérez-Rodríguez et al., 2019). Given the short duration of the study, interannual effects should nevertheless be interpreted with caution, as they may integrate population processes correlated with climatic variability rather than direct effects.

Local management is a consistent structuring factor of tree-level natural enemy occurrence, with heterogeneous response among species. For coccinellids, conventional farming decreased their occurrence, likely reflecting the well-documented negative effects of pesticide use on natural enemies through lethal and sublethal effects (Bommarco et al., 2011; Muneret et al., 2019; Lu et al., 2022; Walker et al., 2025; Tortosa et al., 2025), particularly for coccinellids (Santos et al., 2017; Rasheed et al., 2020; Romero et al., 2025). In contrast, the occurrence of *C. carnea* was higher in conventional orchards, reflecting its relative tolerance (Porcel et al., 2013) and possibly reduced competition and/or intraguild predation with other predators. Despite expected heterogeneity in pesticide use intensity within management categories (Poinas et al., 2026), the contrast between management types in our study remained consistent, indicating a robust local signal beyond variability in individual practices. Nevertheless, a more detailed characterization of farming practices would help disentangle within-category variability. Gradients of management intensity often better explain biodiversity patterns than simple categorical distinctions (Tuck et al., 2014), and integrating such variables at both local and landscape levels could clarify some of the landscape effects detected in our study.

Natural enemy occurrence responded to the surrounding landscape structure. No significant interactions between local management and landscape variables were detected, suggesting mainly additive effects. Model selection identified predictive landscape variables and the spatial scales at which they operated, confirming the scale-dependent influence of the landscape on arthropods (Steffan-Dewenter et al., 2002, Zhang et al., 2020). Landscape configuration rather than composition drove most effects. At the smallest scale (500 m), *C. carnea* benefited from higher aggregation of semi-natural habitats (SNH). In 2022, occurrence increased by as much as threefold along the SNH gradient (Appendix S7), consistent with studies reporting positive effects of SNH availability in agricultural landscapes (Serée et al., 2020; Herrera and Ruano, 2022). At a larger scale (1500 m), coccinellids showed contrasting responses to vineyard configuration, with negative (resp. positive) effects of homogeneous landscapes dominated by large vineyard patches (resp. irregular vineyard patches, i.e. greater edge density). These patterns suggest that landscape homogenization and associated management intensity may constrain sensitive species (Carrié et al., 2017; Stemmelen et al., 2025), while responses remain shaped by distinct aspects of landscape configuration (Zhang et al., 2020). Although responses to landscape structure were evident, connectivity metrics were rarely retained in the final multi-factor models, likely due to strong temporal variation in their effects (Table S3 in Appendix S4). Nevertheless, our results provide empirical support for incorporating multiple habitat types (e.g. SNH) into graph-based models (Savary et al., 2024).

Our results are consistent with previous studies reporting greater arthropod diversity in organic orchards (Weibull et al., 2000; Holzschuh et al., 2007; Caprio et al., 2015; Walker et al., 2025). We also observed interactive effects between local management and landscape structure on diversity (Carrié et al., 2017; Karp et al., 2018; Petit et al., 2017; Ricci et al., 2019). In conventional orchards and at the largest spatial scale, natural enemy evenness increased in landscapes dominated by continuous SNH and decreased in homogeneous landscapes dominated by citrus or vineyard patches. By contrast, evenness remained higher in organic orchards regardless of the surrounding landscape. This result suggests that in conventional orchards, predator assemblages may partly depend on immigration from surrounding SNH, which can provide refuges, overwintering habitats, food resources, and suitable microhabitats (Landis et al., 2000; Bianchi et al., 2006; Tscharntke et al., 2012). In organic orchards, favorable local conditions could potentially buffer predator populations against landscape simplification and management intensification, reducing reliance on spillover from SNH. Interestingly in our system, mealybug infestation severity was lower in organic orchards and in complex natural landscapes (Appendix S2). This pattern supports evidence that SNH enhances biological control (Rusch et al., 2016; Lu et al., 2022), including for citrus mealybugs (Plata et al., 2024). In contrast, leaf miner infestation showed no consistent management effect and variable landscape responses over time.

Our findings offer guidelines to support natural enemies in Corsican citrus orchards, by preserving or restoring semi-natural habitats. Our results also support the benefits of smaller field sizes and higher field edge density in agricultural landscapes (Alignier et al., 2020; Sirami et al., 2020), reinforcing the importance of maintaining heterogeneous crop mosaics to sustain natural enemy communities. Organic management can complement these landscape-based strategies by buffering natural enemy populations against disturbance, contributing to more stable pest suppression over time, although responses vary among species. The species-specific responses documented here can inform ecological models by providing parameters relating predator presence and abundance to environmental and landscape variables. From a methodological standpoint, future studies examining similar biological responses should focus on multiple sites and years rather than intensive sampling of a large number of trees across a few plots. Future surveys should also integrate functional measures of predation, such as sentinel prey experiments or direct measurement of predation rates, to better link predator density and diversity with pest control services (Masson et al., 2025; Lavigne et al., 2025).

In summary, this research indicates that combined local and landscape management approaches, particularly organic farming and the preservation of semi-natural habitats, can support natural enemies and their potential to reduce pest pressure in Corsican citrus orchards. Long-term ecological monitoring will be critical to refine these strategies and to extend findings to other Mediterranean perennial cropping systems.

## AUTHOR CONTRIBUTIONS

L.J., E.C^1^., E.C^4^., M.P., S.S., V.R. conceived and designed the research. L.J, L.M. and E.C^2^. contributed to the sampling design and collected field data. E.F. and E.C^1^. carried out the land cover processing. E.C^1^. analysed the data and wrote the first version of the manuscript. All authors critically revised the manuscript and gave final approval for publication.

## ACKNOWLEDGEMENTS

L.M. was funded by the LIFE project ‘Vida for Citrus’ (LIFE18 CCA/ES/001109). The work of E.C^1^. and E.F. was supported by the French ECOPHYTO II+/OFB projects IMPACT (convention n° 23-1315) and IRIS (convention n° 23-1316). We warmly thank all citrus growers, as well as their grower associations, for granting access to their orchards and for their contribution to this study. We also warmly and sincerely thank AOP Fruits de Corse for their invaluable collaboration and strong support throughout this study. We are particularly grateful to Camille Portes and Marine Marjou (INRAE, BioSP) for their essential role in facilitating access to data and code.

## DATA AVAILABILITY STATEMENT

The natural enemy data used in this study will be made available on request, upon manuscript acceptance. The soil and topographic data are openly available via https://doi.org/10.1017/eds.2024.50 (Portes et al., 2024). Climatic data are available on request from the SICLIMA platform (https://agroclim.inrae.fr/siclima/).

