## Supplemental information for "Scouting ecological drivers of natural enemies in citrus orchards: implications for biological control in the Corsican agricultural landscape"

Appendix S1. Spatial design of arthropod sampling

Appendix S2. Pest infestation levels

Appendix S3. Climate, soil and topographic explanatory variables

Appendix S4. Functional connectivity analysis

Appendix S5. Additional cross-validation analyses

Appendix S6. Relationships between site-level natural enemy diversity and evenness, and landscape structure across spatial scales

Appendix S7. Effects of surrounding semi-natural habitats on *Chrysoperla carnea* occurrence

#### Appendix S1 - Spatial design of arthropod sampling

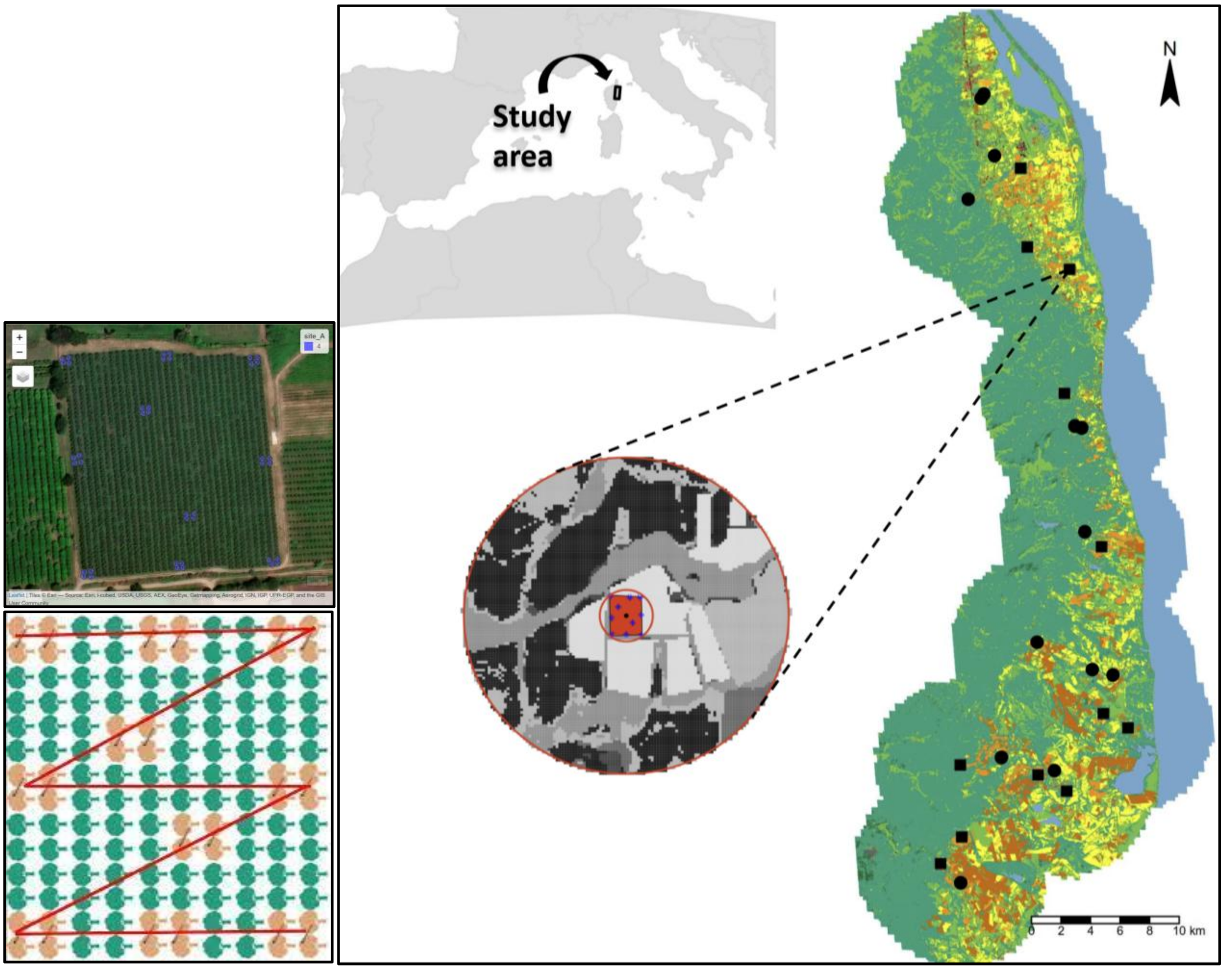


**FIGURE S1.** Study area and sampling design. Locations of clementine orchards selected for this study in eastern Corsica are shown. Insets illustrate the regional context of the study area, the spatial distribution of sampled orchards across the landscape, and the local sampling design within orchards. Within each orchard, arthropods were surveyed on 40 trees following a standardized back-and-forth transect comprising 10 evenly spaced sampling stops, with four adjacent trees sampled at each stop. Landscape structure was quantified at multiple spatial scales (500, 1500, and 3000 m buffers from the orchard edge); the 500-m radius buffer is shown here for illustration.

### Appendix S2 - Pest infestation levels

The effects of temporal, management and landscape factors on infestation severity of vegetative flushes by mealybugs (*Planococcus citri*) and leafminers (*Phyllocnistis citrella*) were examined using Linear Mixed Models (LMMs), followed by type-III tests of deviance (or type-II when no interaction was included). The analysis was conducted at the tree scale, including only trees with at least one infested flush. The response variable, defined as the proportion of infested flushes relative to the total number of flushes, was log-transformed to meet normality assumptions of model residuals. In all models, orchard identity was included as a random effect. A first analysis included time of the year (season x year), local management and their two-way interaction as fixed effects. A second analysis tested the effects of landscape (GYRATE_CV, 3000 m), management and their two-way interactions as fixed effects. Finally, a third analysis tested whether landscape effects varied across time by including time of the year, landscape and their two-way interactions as fixed effects.

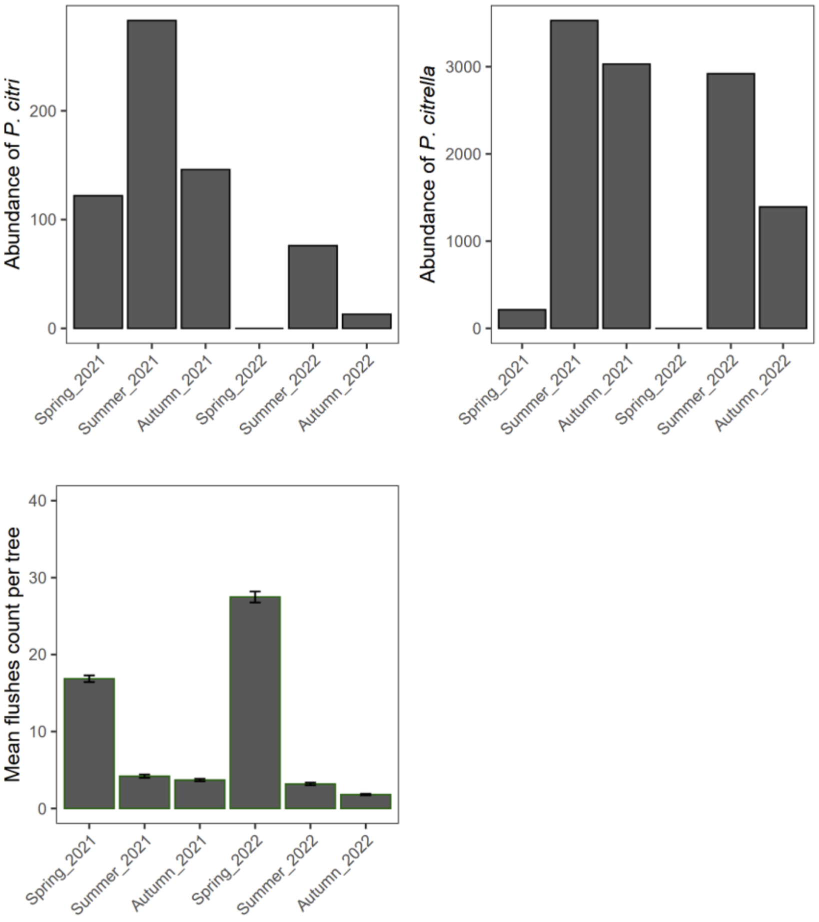


**FIGURE S2.** Seasonal abundance of *P. citri* (left) and *P. citrella* (right) across sampled clementines orchards (n = 25), and mean number of vegetative flushes per tree (bottom panel) for the sampling periods (spring, summer and autumn of 2021 and 2022). Bars represent total observed abundances (top panels) and mean flush counts per tree ± SE (bottom panel).

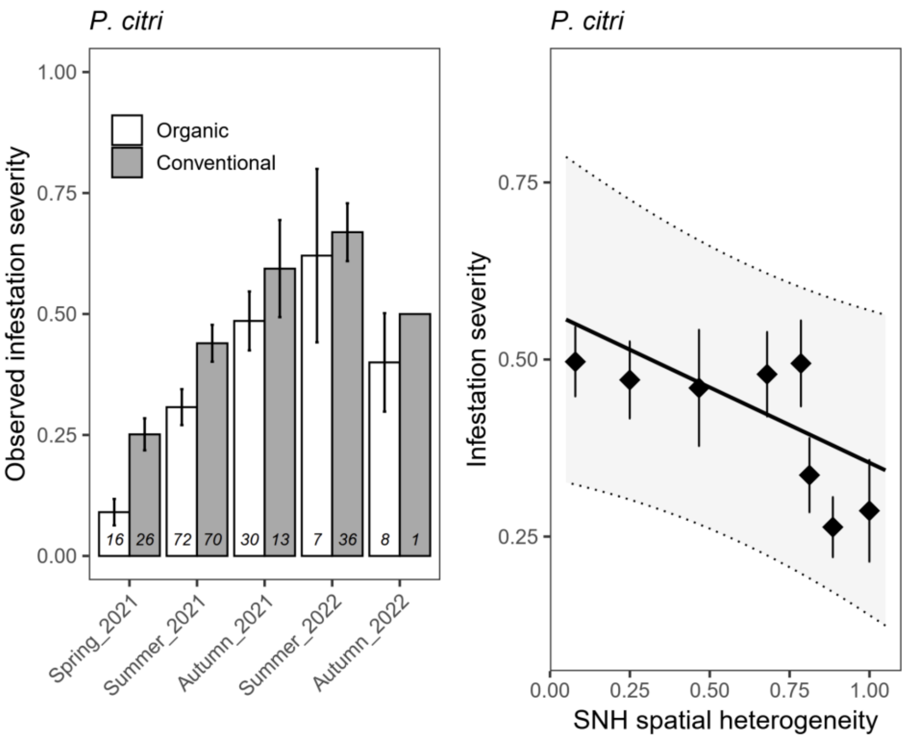


**FIGURE S3.** Observed mean infestation severity (± SE) of mealybug according to the time of the year and the management systems (organic *vs* conventional) (left). The number of trees per condition is given at the bottom of each bar. Mealybug infestation severity according to the surrounding SNH spatial heterogeneity (GYRATE_CV, 3000 m), across both management systems (right). Solid and dotted lines represent predictions and their 95% confidence intervals. Observed data were split into groups of homogeneous size (~279:9). Points and segments correspond to observed class means of equal sizes ± SE.

**Mealybugs infestation**

The proportion of infested flushes by mealybugs was significantly affected by the time of the year ($P < 0.001$) and local management ($P < 0.05$). Infestation severity increased from spring 2021 (0.10c) to summer 2021 (0.27b) until autumn 2021 (0.54a), summer 2022 (0.44a) and autumn 2022 (0.41ab). No infestation occurred in spring 2022 (Figure S2). It was on average higher in conventional (0.36) than in organic orchards (0.25). SNH spatial heterogeneity (GYRATE_CV, 3000 m), had a significant negative effect on infestation severity (Figure S3 right panel, $P < 0.05$). These effects were consistent across management systems and the studied periods, as no significant interaction was detected (Table S1).

**Leafminers infestation**

The proportion of infested flushes by leafminers was significantly affected by the interaction between time of the year and management ($P < 0.001$, Table S1). In spring 2021, infestation severity was low and similar in conventional (0.16d) and organic orchards (0.14d). Infestation severity increased sharply thereafter, with no significant difference between the two management types (conventional *vs* organic) per period: summer 2021 (0.85ab *vs* 0.84bc), autumn 2021 (0.76c *vs* 0.83bc), summer 2022 (0.93ab *vs* 0.99a), autumn 2022 (0.90ab *vs* 0.92bc). No significant interactions were found between the landscape variable and local management (Table S1), but the interaction between time of the year and the landscape variable was significant ($P < 0.001$, Table S1). SNH spatial heterogeneity negatively affected infestation in spring 2021 (slope = -0.53, 95% CI [-0.81, -0.24]) and autumn 2022 (slope = -0.26, 95% CI [-0.44, -0.09]), while slopes were not significant in other periods.

**TABLE S1:** Summary of linear mixed models testing the effects of temporal, local management (organic *vs* conventional) and landscape factors, namely the spatial heterogeneity of semi-natural habitats (SNH GYRATE_CV within 3000 m buffers) on severity of mealybug and leafminers infestation. Only significance levels are shown for clarity. Arrows indicate fixed effects included in each model. Two-way interaction terms are reported on separate rows. Significance level was set at $P = 0.05$.

| | **Mealybugs** | **Leafminers** |
|---|---|---|
| **Model 1:** tests temporal and management effects | | |
| ➢ Time | ***P* < 0.001** | ***P* < 0.001** |
| ➢ Management | ***P* < 0.05** | *ns* |
| Time : Management | *ns* | ***P* < 0.001** |
| **Model 2:** tests landscape and management effects | | |
| ➢ SNH GYRATE_CV | ***P* < 0.05** | *ns* |
| ➢ Management | *ns* | *ns* |
| SNH GYRATE_CV : Management | *ns* | *ns* |
| **Model 3:** tests landscape and temporal effects | | |
| ➢ SNH GYRATE_CV | ***P* < 0.05** | ***P* < 0.001** |
| ➢ Time | ***P* < 0.001** | ***P* < 0.001** |
| SNH GYRATE_CV : Time | *ns* | ***P* < 0.001** |

#### Appendix S3 - Climate, soil and topographic explanatory variables

**TABLE S2:** Climate, soil and topographic variables used in the analyses, their definitions, temporal aggregation applied and data sources.

| Name | Full definition | Temporal integration | Source |
|---|---|---|---|
| BIO1 | Annual mean air temperature (°C) | Yearly | https://pubs.usgs.gov/publication/ds691 |
| BIO2 | Annual mean diurnal range (°C) | Yearly | https://pubs.usgs.gov/publication/ds691 |
| BIO3 | Isothermality (%): (BIO2/BIO7) * 100 | Yearly | https://pubs.usgs.gov/publication/ds691 |
| BIO4 | Temperature seasonality (SD, °C) | Yearly | https://pubs.usgs.gov/publication/ds691 |
| BIO5 | Max temperature of warmest month (°C) | Yearly | https://pubs.usgs.gov/publication/ds691 |
| BIO6 | Min temperature of coldest month (°C) | Yearly | https://pubs.usgs.gov/publication/ds691 |
| BIO7 | Annual temperature range (°C) | Yearly | https://pubs.usgs.gov/publication/ds691 |
| BIO8 | Mean temperature of wettest quarter (°C) | Yearly | https://pubs.usgs.gov/publication/ds691 |
| BIO9 | Mean temperature of driest quarter (°C) | Yearly | https://pubs.usgs.gov/publication/ds691 |
| BIO10 | Mean temperature of warmest quarter (°C) | Yearly | https://pubs.usgs.gov/publication/ds691 |
| BIO11 | Mean temperature of coldest quarter (°C) | Yearly | https://pubs.usgs.gov/publication/ds691 |
| BIO12 | Annual precipitation (mm) | Yearly | https://pubs.usgs.gov/publication/ds691 |
| BIO13 | Precipitation of wettest month (mm) | Yearly | https://pubs.usgs.gov/publication/ds691 |
| BIO14 | Precipitation of driest month (mm) | Yearly | https://pubs.usgs.gov/publication/ds691 |
| BIO15 | Precipitation seasonality (CV, %) | Yearly | https://pubs.usgs.gov/publication/ds691 |
| BIO16 | Precipitation of wettest quarter (mm) | Yearly | https://pubs.usgs.gov/publication/ds691 |
| BIO17 | Precipitation of driest quarter (mm) | Yearly | https://pubs.usgs.gov/publication/ds691 |
| BIO18 | Precipitation of warmest quarter (mm) | Yearly | https://pubs.usgs.gov/publication/ds691 |
| BIO19 | Precipitation of coldest quarter (mm) | Yearly | https://pubs.usgs.gov/publication/ds691 |
| ff_q | Daily wind speed at 10 m (m/s) | Seasonal | https://agroclim.inrae.fr/siclima/help/decouvrir/extraction.html |
| t_q | Daily mean air temperature (°C) | Seasonal | https://agroclim.inrae.fr/siclima/help/decouvrir/extraction.html |
| tsup_h_q | Daily maximum air temperature (°C) | Seasonal | https://agroclim.inrae.fr/siclima/help/decouvrir/extraction.html |
| tinf_h_q | Daily minimum air temperature (°C) | Seasonal | https://agroclim.inrae.fr/siclima/help/decouvrir/extraction.html |

| | | | |
|---|---|---|---|
| runc_q | Daily runoff (mm) | Seasonal | https://agroclim.inrae.fr/siclima/help/decouvrir/extraction.html |
| ssi_q | Daily incident shortwave radiation ($J/cm^2$) | Seasonal | https://agroclim.inrae.fr/siclima/help/decouvrir/extraction.html |
| dli_q | Daily incident longwave radiation ($J/cm^2$) | Seasonal | https://agroclim.inrae.fr/siclima/help/decouvrir/extraction.html |
| prenei_q | Daily solid precipitation (mm) | Seasonal | https://agroclim.inrae.fr/siclima/help/decouvrir/extraction.html |
| preliq_q | Daily liquid precipitation (mm) | Seasonal | https://agroclim.inrae.fr/siclima/help/decouvrir/extraction.html |
| swi_q | Daily soil moisture index (%) | Seasonal | https://agroclim.inrae.fr/siclima/help/decouvrir/extraction.html |
| q_q | Daily specific humidity (g/kg) | Seasonal | https://agroclim.inrae.fr/siclima/help/decouvrir/extraction.html |
| hu_q | Daily relative humidity (%) | Seasonal | https://agroclim.inrae.fr/siclima/help/decouvrir/extraction.html |
| evap_q | Daily actual evapotranspiration (mm) | Seasonal | https://agroclim.inrae.fr/siclima/help/decouvrir/extraction.html |
| etp_q | Daily potential evapotranspiration (mm) | Seasonal | https://agroclim.inrae.fr/siclima/help/decouvrir/extraction.html |
| Soil chemical elements | Clay content; Fine silt content; Coarse silt content; Fine sand content; Coarse sand content; Residual water content; Water pH; Exchangeable aluminum; Total aluminum; Boron soluble in boiling water; Organic carbon; Exchangeable calcium; Total calcium; Total limestone; Cation exchange capacity (CEC); Exchangeable iron; Free iron; Total iron; Exchangeable potassium; Total potassium; Organic matter; Exchangeable magnesium; Total magnesium; Exchangeable manganese; Total manganese; Total nitrogen; Exchangeable sodium; Total sodium; Available phosphorus; Total arsenic; Extractable cadmium; Total cadmium; Total cobalt; Extractable chromium; Extractable copper; Total copper; Total mercury; Total molybdenum; Extractable nickel; Total nickel; Extractable lead; Total lead; Total thallium; Extractable zinc; Total zinc | - | https://doi.org/10.1017/eds.2024.50 |
| Relief and orientation | Altitude; Proportion of the cell corresponding to the cardinal (N, E, S, W) and intercardinal (N-E, S-E, S-W, N-W) directions | - | https://doi.org/10.1017/eds.2024.50 |

#### Appendix S4 - Functional connectivity analysis

Landscape graphs were constructed to model species-specific ecological networks. Habitat patches were defined according to species ecology and represented as nodes, while links corresponded to least-cost paths computed from a resistance surface. Five resistance classes were defined based on species' ability to cross and persist within landscape elements: (1) habitat, (2) favourable, (3) neutral, (4) unfavourable and (5) ecological barriers. Resistance values ranged from 1 (habitat) to 1000 (barrier), with intermediate values defined either logarithmically (684, 900, 968) or exponentially (6, 32, 178) (Table S2).

Maximum dispersal distances (Dmax) used in the graph model (Table S2) correspond to short-range functional dispersal relevant to seasonal dynamics rather than long-distance migratory capacity. Least-cost distances were computed in metric units and converted to cost units using a linear relationship between Euclidean and cost distances.

**TABLE S3:** Species ecological parameters (maximum dispersal capacity $D_{max}$, minimum surface area $S_{min}$) and resistance values attributed according to the five categories: 'habitat', 'favorable', 'neutral' , 'unfavorable', 'ecological barrier', across the land class: (1) urban areas (dense and dispersed), (2) industrial and commercial zones, (3) roads, (4) oilseed crops, (5) cereals, (6) legumes, (7) soybean, (8) sunflower, (9) corn, (10) rice, (11) tubers, (12) cultivated grasslands, (13) orchards, (14) vineyards, (15) semi-natural habitats (deciduous forest, coniferous forest and heathlands), (16) artificial grasslands, (17) minerals, (18) sands, (19) snow, (20) water, (21) citrus orchards.

| Species | | | *Chrysoperla carnea* | *Exochomus quadripustulatus* | *Cryptolaemus montrouzieri* | *Harmonia axyridis* |
|---|---|---|---|---|---|---|
| $D_{max}$ (m) | | | 1500 | 500 | 750 | 1500 |
| $S_{min}$ (ha) | | | 0.01 | 0.01 | 0.01 | 0.01 |
| | *Resistances values* | | | | | |
| Land class | 1<br>6<br>32<br>178<br>1000 | 1<br>684<br>900<br>968<br>1000 | 21<br>13,14,15,<br>4,6,7,8,9,10,11,12,16<br>1,2,3,17<br>18,19,20 | 21<br>13,14,15,<br>4,6,7,8,9,10,11,12,16<br>1,2,3,17<br>18,19,20 | 21<br>13,14,15,<br>1,4,6,7,8,9,10,11,12,16<br>2,3,17<br>18,19,20 | 21<br>1,12,13,14,15,<br>4,6,7,8,9,10,11,16<br>2,3,17<br>18,19,20 |
| References | | | Duelli, 1980<br>Paredes et al., 2024 | Van der Werf et al., 2000 | Maes et al., 2014 | Osawa, 2000<br>Koch, 2003<br>Maes et al., 2014 |

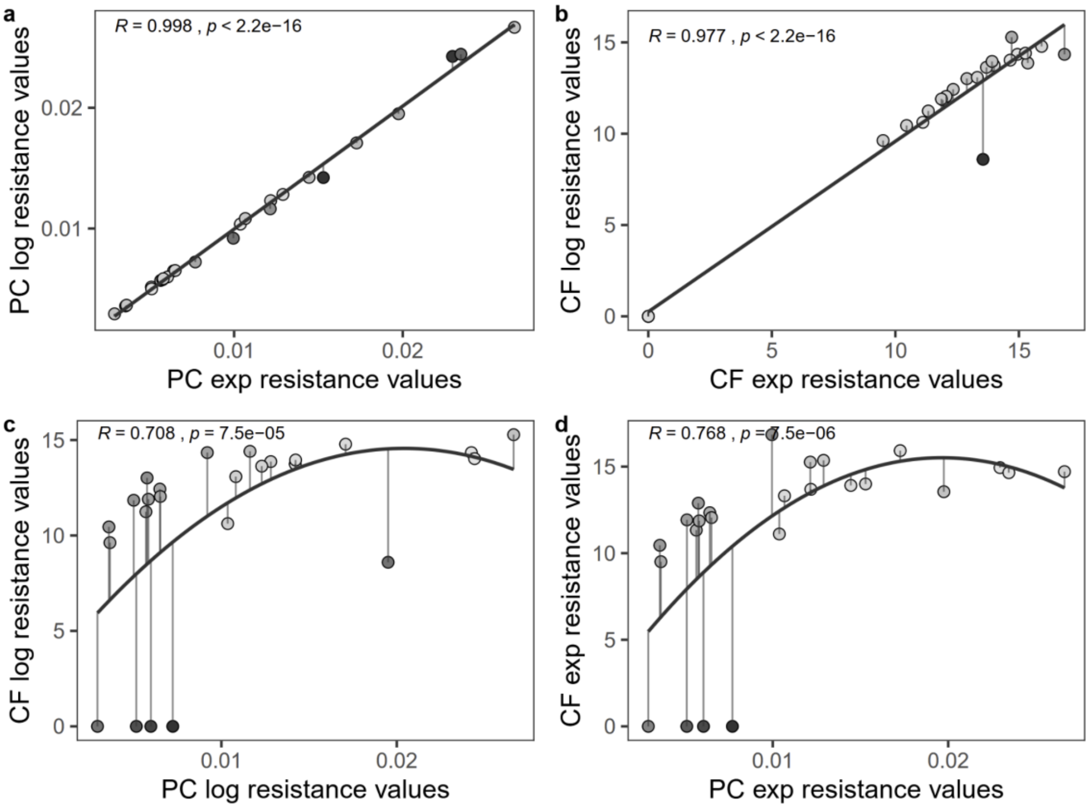


**FIGURE S4.** Relationships between connectivity metrics calculated using the R package ‘graph4lg’ according to the resistance transformation (logarithmic or exponential) for PC (a) and CF (b). Panels (c) and (d) show relationships between PC and CF calculated using logarithmic (c) or exponential (d) resistance gradients. PC and CF values are transformed for visualization purposes.Vertical segments and point colour gradients represent the deviation between predicted and observed values along the y-axis. Pearson correlation coefficients are reported for linear relationships (panels a and b), whereas Spearman rank correlations are shown for non-linear relationships (panels c and d).

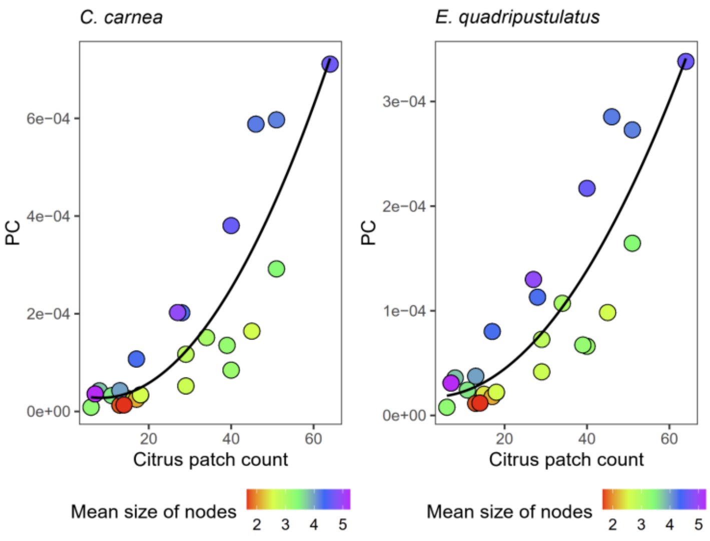


**FIGURE S5.** Relationship between the Probability of Connectivity (PC) and the number of citrus patches within a 3 km buffer around each orchard for illustration purposes (*C.carnea*; left; *E. quadripustulatus* right). Point colors indicate mean size of nodes (citrus patches) in the buffer.

**TABLE S4:** Summary of generalised linear mixed models (GLMM) testing the effects of the season, local management and connectivity metrics (PC and CF), and their two-way interactions, on the probability of predator occurrence at the tree scale. Arrows indicate fixed effects included in each model. Significant interactions (in bold) are illustrated in Fig. S6.

| ***Species*** | *Chrysoperla carnea* | | *Exochomus quadripustulatus* | | *Cryptolaemus montrouzieri* | | *Harmonia axyridis* | |
|---|---|---|---|---|---|---|---|---|
| ***Year*** | ***2021*** | ***2022*** | ***2021*** | ***2022*** | ***2021*** | ***2022*** | ***2021*** | ***2022*** |
| **Model 1:** tests seasonal, local management and PC effects | | | | | | | | |
| ➢ Season | P<0.05 | P<0.001 | *ns* | *ns* | P<0.001 | P<0.001 | P<0.001 | P<0.001 |
| ➢ Management | P<0.05 | P<0.05 | P<0.001 | *ns* | P<0.05 | P<0.001 | *ns* | *ns* |
| ➢ PC | *ns* | *ns* | P<0.05 | *ns* | *ns* | *ns* | *ns* | *ns* |
| Season : PC | **P<0.01** | *ns* | **P<0.01** | *ns* | **P<0.05** | *ns* | **P<0.05** | *ns* |
| Management : PC | *ns* | *ns* | *ns* | *ns* | 0.05. | *ns* | *ns* | *ns* |
| **Model 2:** tests seasonal, local management and CF effects | | | | | | | | |
| ➢ Season | *ns* | P<0.001 | P<0.001 | *ns* | P<0.001 | P<0.001 | P<0.001 | P<0.001 |
| ➢ Management | P<0.001 | P<0.05 | P<0.001 | *ns* | P<0.05 | P<0.001 | 0.09 . | *ns* |
| ➢ CF | 0.07 . | *ns* | *ns* | *ns* | *ns* | 0.06. | *ns* | *ns* |
| Season : CF | 0.06 . | *ns* | *ns* | *ns* | **P<0.05** | *ns* | **P<0.01** | *ns* |
| Management : CF | **P<0.01** | *ns* | *ns* | *ns* | *ns* | *ns* | *ns* | *ns* |

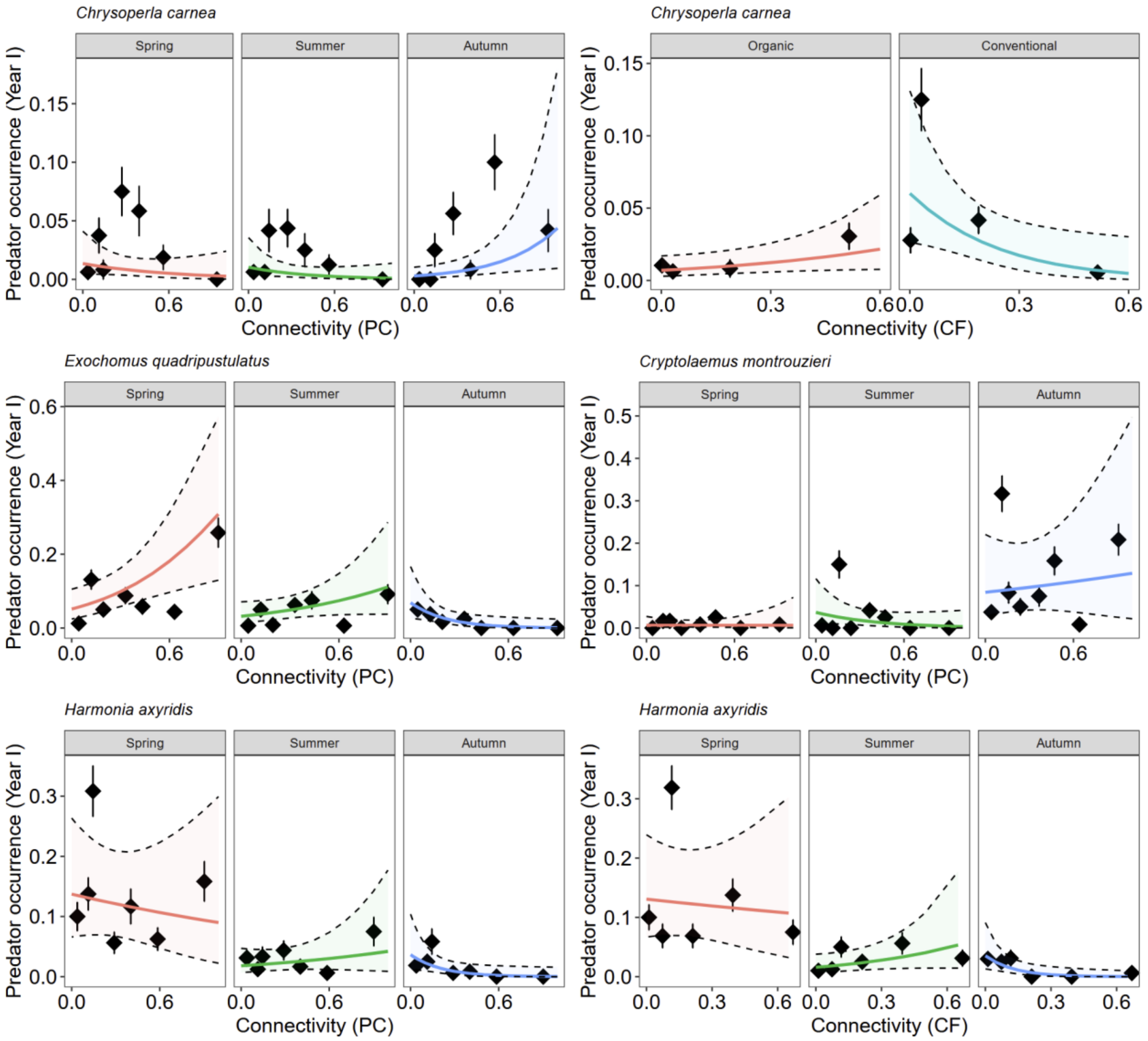


**FIGURE S6.** Predator occurrence at the tree scale as a function of experimental factors (year, season, local management) and functional connectivity metrics (PC and CF) for *C. carnea* (top panels), *E. quadripustulatus* and *C. montrouzieri* (middle panels), and *H. axyridis* (bottom panels). Solid lines represent GLMM-predicted values (± 95% confidence intervals). Points and segments correspond to observed class means of equal sizes ± SE. PC and CF values were min–max scaled to range between 0 and 1 prior to analysis.

#### Appendix S5 - Additional cross-validation analyses

To further illustrate the robustness of model performance, we implemented an additional site-level cross-validation, in which all observations from a given site were iteratively excluded from model training and used exclusively for testing.

Models were fitted using a fixed sampling intensity corresponding to the maximum number of trees per site (n = 40). Site-level random effects were excluded to ensure that predictions relied solely on fixed effects.

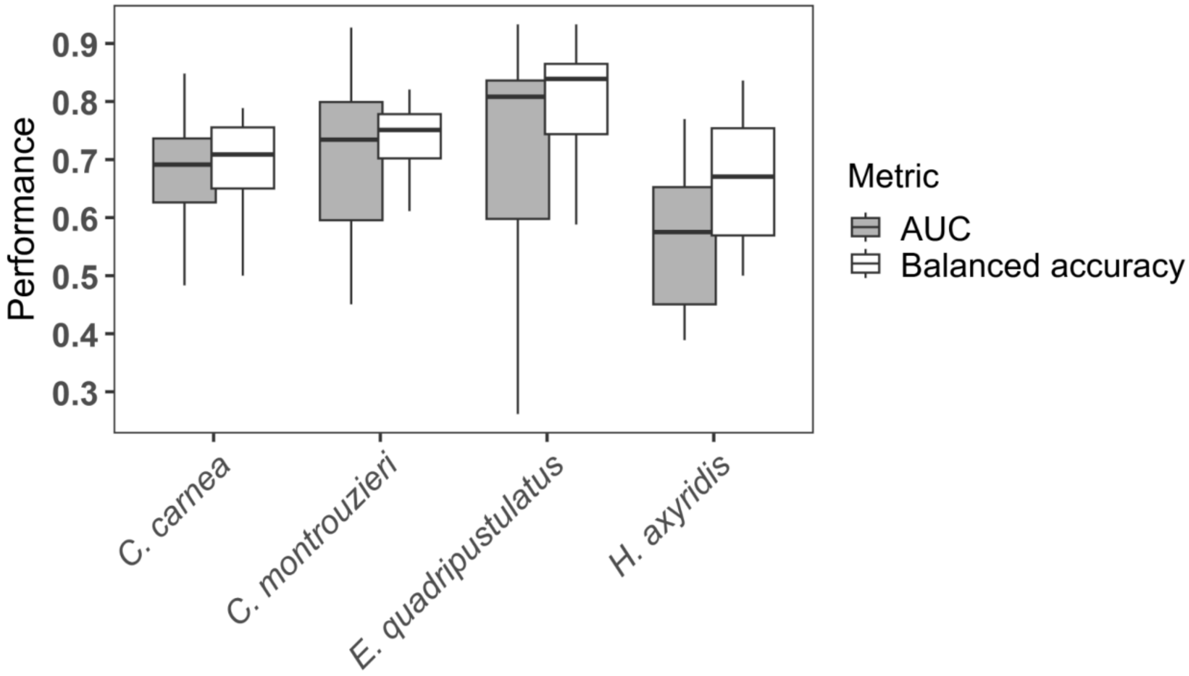


**FIGURE S7.** Model performance under leave-one-site-out cross-validation. For each species, final environmental GLMMs were trained on all sites except one and evaluated on the corresponding held-out site. Performance metrics include area under the ROC curve (AUC) and balanced accuracy. Boxplots represent the distribution of scores across validation runs (750 iterations).

### Appendix S6 - Relationships between site-level natural enemy diversity and evenness, and landscape structure across spatial scales

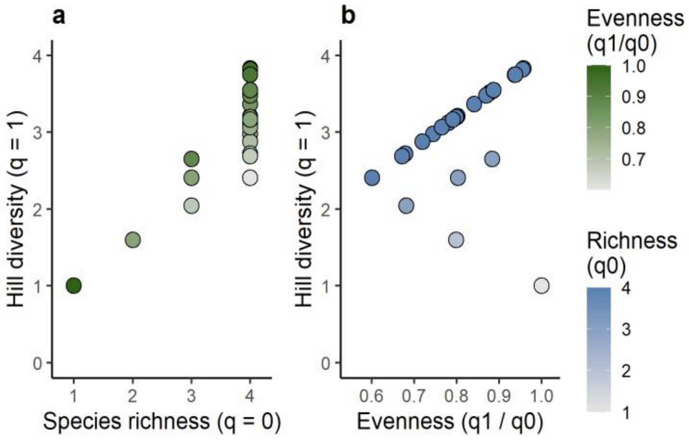


**FIGURE S8.** Relationship between species richness (q = 0) and Hill diversity (q = 1), showing that diversity varies substantially within each richness level (a). Relationship between Hill diversity (q = 1) and evenness (q1/q0) (b). Together, these patterns indicate that, given the limited number of species, variation in Hill diversity (q = 1) primarily reflects differences in species' occurrence frequencies (i.e. evenness) rather than differences in richness.

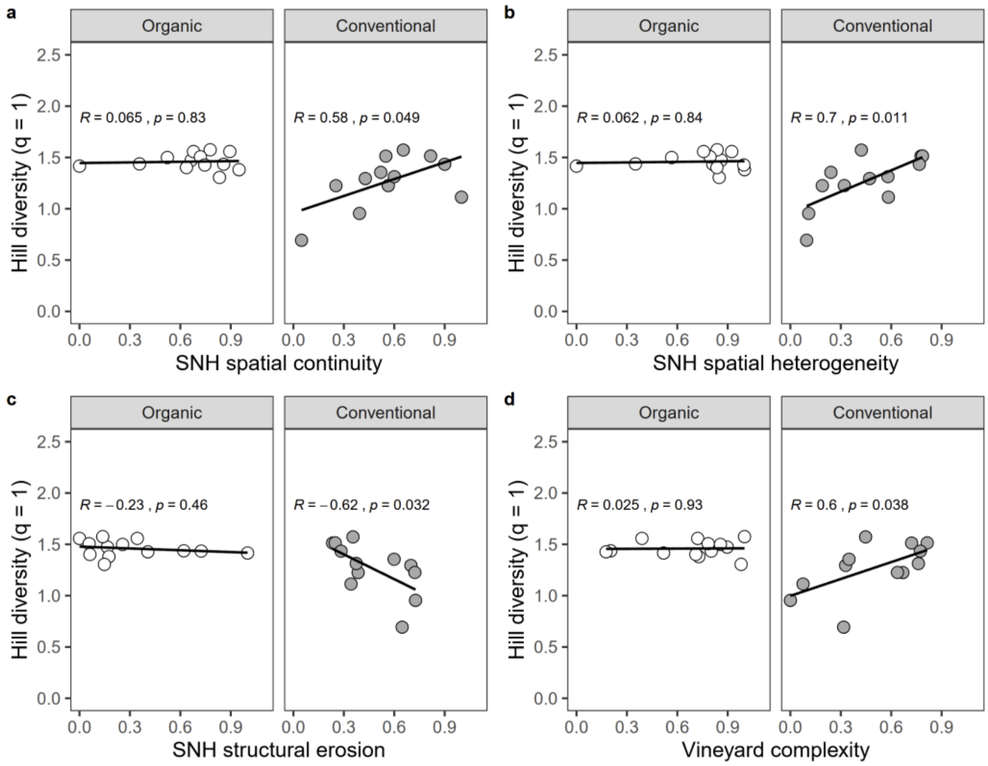


**FIGURE S9.** Relationships between site-level natural enemy evenness (log-transformed Hill diversity, q = 1) and landscape metrics at the 3 km scale, according to the local management type. Panels show the effects of class-level metrics: (a) SNH spatial continuity (COHESION), (b) SNH spatial heterogeneity (GYRATE_CV), (c) SNH structural erosion (FRAC_MN), and (d) vineyard structural complexity (PAFRAC). Lines represent linear fits, with Pearson correlation coefficients (*R*) and associated *P*-values indicated in each panel.

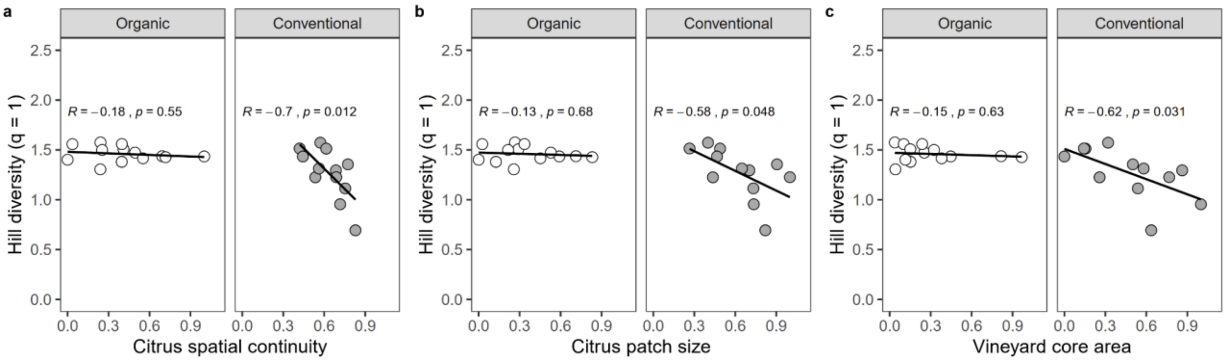


**FIGURE S10.** Relationships between site-level natural enemy evenness (log-transformed Hill diversity, q = 1) and landscape metrics at the 3 km scale, according to the local management type. Panels show the effects of class-level metrics: (a) Citrus spatial continuity (COHESION), (b) Citrus patch size (AREA_MN), and (c) vineyard core area (CORE_MN). Lines represent linear fits, with Pearson correlation coefficients (*R*) and associated *P*-values indicated in each panel.

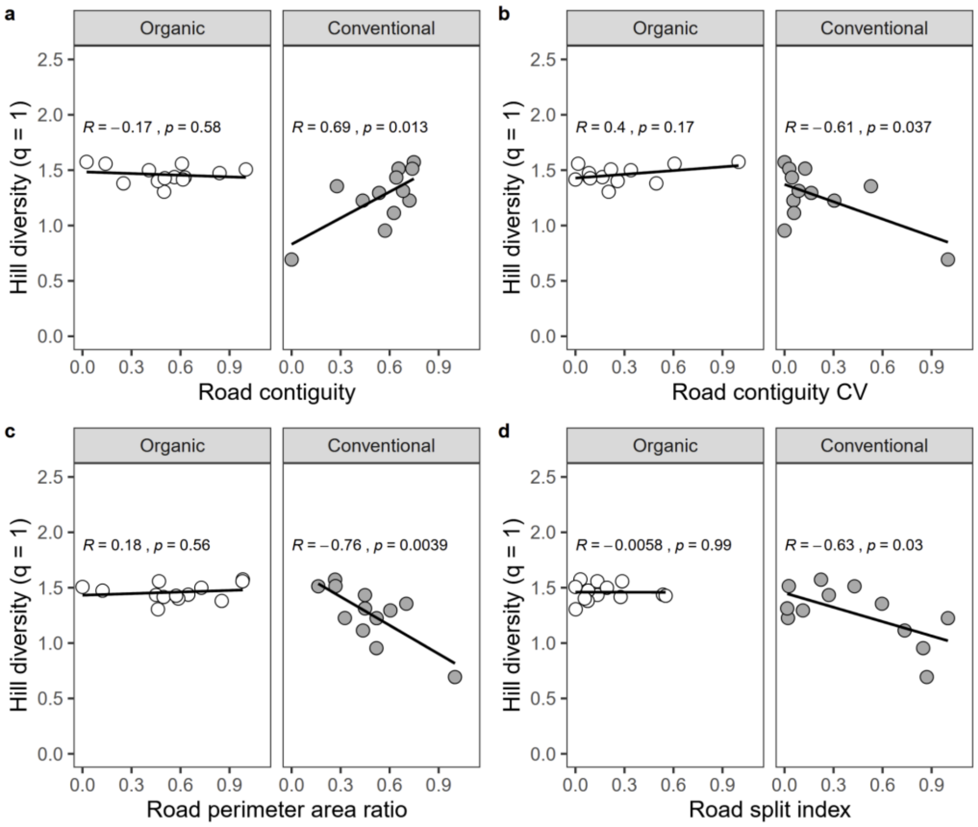


**FIGURE S11.** Relationships between site-level natural enemy evenness (log-transformed Hill diversity, q = 1) and landscape metrics at the 500-meter scale, according to the local management type. Panels show the effects of class-level metrics: (a) Road contiguity (CONTIG_MN), (b) Road contiguity coefficient of variation, CV (CONTIG_CV), (c) Road perimeter to area ratio (PARA_MN), and (d) Road split index (SPLIT). Lines represent linear fits, with Pearson correlation coefficients (*R*) and associated *P*-values indicated in each panel.

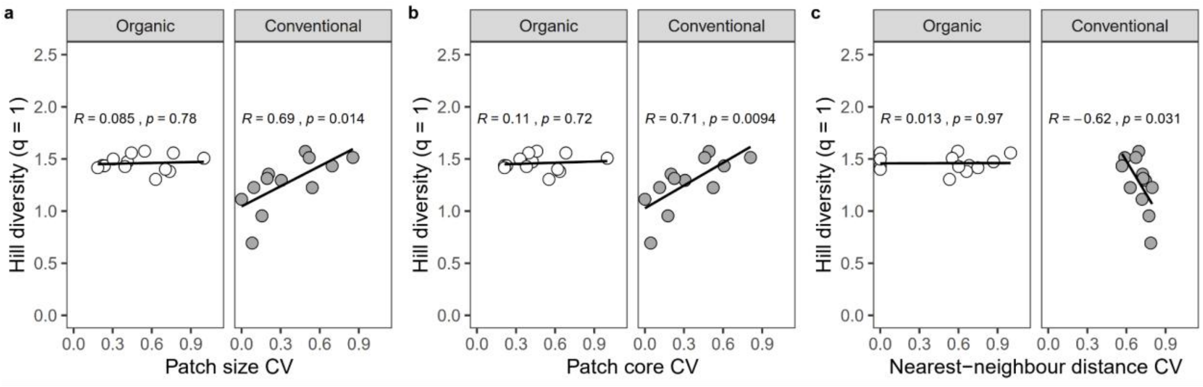


**FIGURE S12.** Relationships between site-level natural enemy evenness (log-transformed Hill diversity, q = 1) and landscape metrics at the 3 km scale, according to the local management type. Panels show the effects of landscape-level metrics: (a) Patch size coefficient of variation, CV (AREA_CV), (b) Patch core CV (CORE_CV), and (c) Nearest-neighbour distance CV (ENN_CV). Lines represent linear fits, with Pearson correlation coefficients (*R*) and associated *P*-values indicated in each panel.

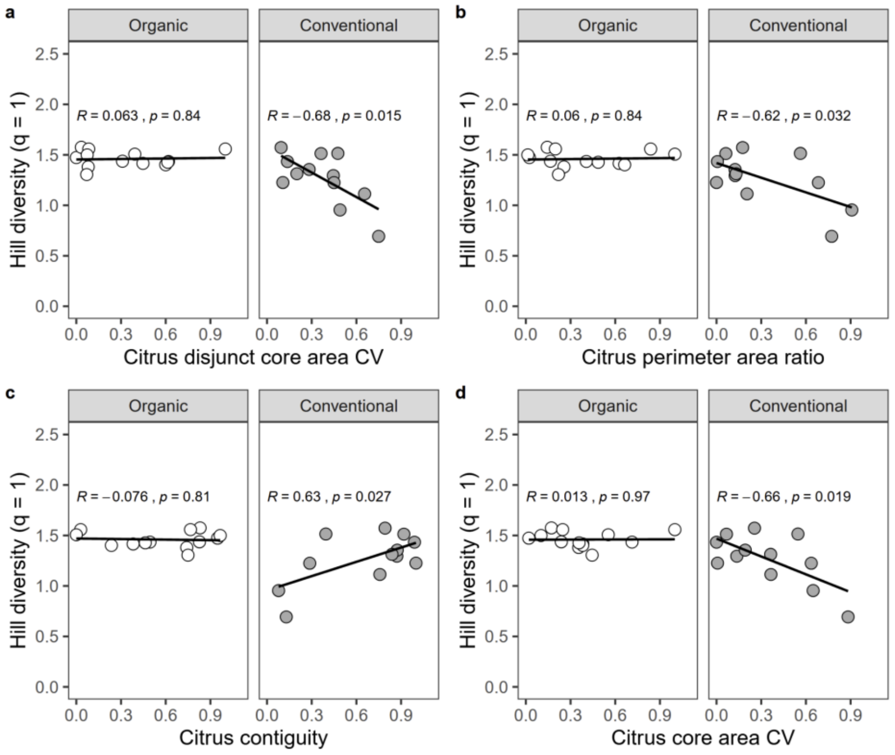


**FIGURE S13.** Relationships between site-level natural enemy evenness (log-transformed Hill diversity, q = 1) and landscape metrics at the 1.5 km scale, according to the local management type. Panels show the effects of class-level metrics: (a) Citrus disjunct core area coefficient of variation, CV (DCORE_CV), (b) Citrus perimeter to area ratio (PARA_MN), (c) Citrus contiguity (CONTIG_MN), and (d) Citrus core area CV (CORE_CV). Lines represent linear fits, with Pearson correlation coefficients (*R*) and associated *P*-values indicated in each panel.

### Appendix S7 - Effects of surrounding semi-natural habitats on *Chrysoperla carnea* occurrence.

**TABLE S5:** Summary of generalized linear mixed models (GLMMs) testing, within each survey year, the effects of season, local management (organic *vs*. conventional), and a landscape variable, namely the aggregation of semi-natural habitats (SNH AI within 500 m buffers), and their two-way interactions on the probability of *Chrysoperla carnea* occurrence at the tree scale. Only significance levels are shown for clarity. Arrows indicate fixed effects included in each model. Significant landscape effects (in bold) are illustrated in Figure S14.

| ***Year*** | ***2021*** | ***2022*** |
|---|---|---|
| ➢ Season | P < 0.01 | P < 0.001 |
| ➢ Management | P < 0.05 | P < 0.05 |
| ➢ SNH AI 0.5 | *ns* | **P < 0.001** |
| Season : SNH AI 0.5 | **P < 0.001** | *ns* |
| Management : SNH AI 0.5 | *ns* | *ns* |

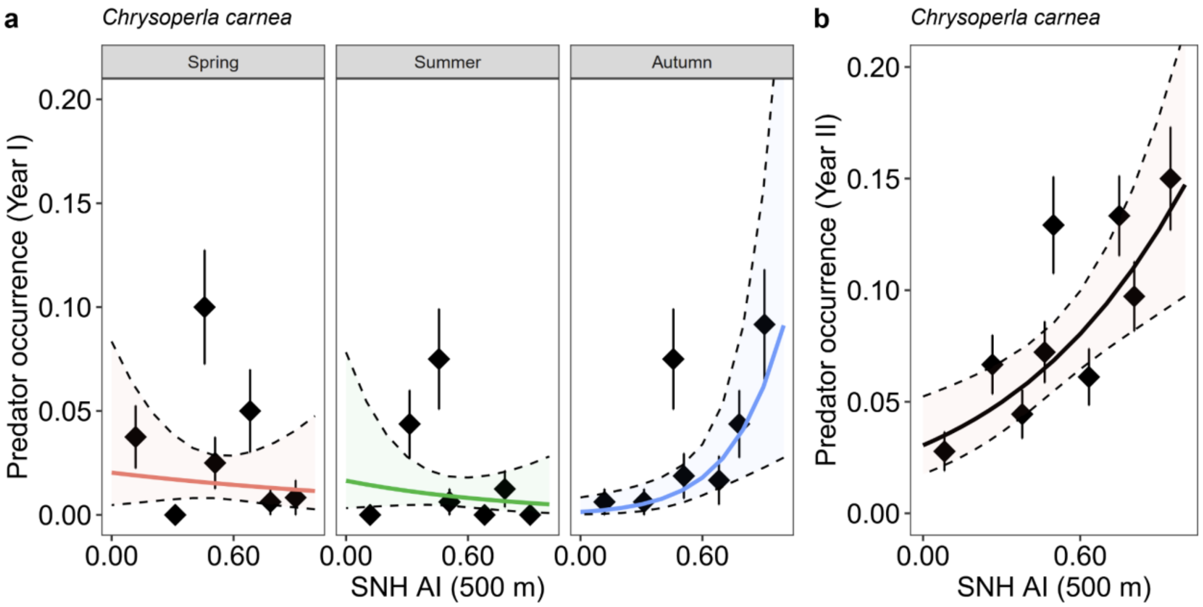


**FIGURE S14.** Predator occurrence of *Chrysoperla carnea* at the tree scale in relation to the aggregation index of semi-natural habitats (SNH AI, within 500 m buffers). In the first year (panel a), occurrence is shown according to season, whereas in the second year (panel b), it is shown as a function of SNH AI only. Solid lines represent GLMM-predicted values, with shaded areas indicating 95% confidence intervals. Points and segments correspond to observed class means of equal sizes ± SE.
